# Mitoregulin supports mitochondrial membrane integrity and protects against cardiac ischemia-reperfusion injury

**DOI:** 10.1101/2024.05.31.596875

**Authors:** Colleen S. Stein, Xiaoming Zhang, Nathan H. Witmer, Edward Ross Pennington, Saame Raza Shaikh, Ryan L. Boudreau

**Author notes:** To whom correspondence should be addressed: Ryan L. Boudreau 4334 PBDB, University of Iowa, Department of Internal Medicine Iowa City, IA 52242.

## Abstract

We and others discovered a highly-conserved mitochondrial transmembrane microprotein, named Mitoregulin (Mtln), that supports lipid metabolism. We reported that Mtln strongly binds cardiolipin (CL), increases mitochondrial respiration and Ca^2+^ retention capacities, and reduces reactive oxygen species (ROS). Here we extend our observation of Mtln-CL binding and examine Mtln influence on cristae structure and mitochondrial membrane integrity during stress. We demonstrate that mitochondria from constitutive- and inducible Mtln-knockout (KO) mice are susceptible to membrane freeze-damage and that this can be rescued by acute Mtln re-expression. In mitochondrial-simulated lipid monolayers, we show that synthetic Mtln decreases lipid packing and monolayer elasticity. Lipidomics revealed that Mtln-KO heart tissues show broad decreases in 22:6-containing lipids and increased cardiolipin damage/remodeling. Lastly, we demonstrate that Mtln-KO mice suffer worse myocardial ischemia-reperfusion injury, hinting at a translationally-relevant role for Mtln in cardioprotection. Our work supports a model in which Mtln binds cardiolipin and stabilizes mitochondrial membranes to broadly influence diverse mitochondrial functions, including lipid metabolism, while also protecting against stress.

## INTRODUCTION

Microproteins have emerged as a class of proteins often found to be encoded by misannotated long noncoding RNAs (lncRNAs). We and others previously discovered that a ubiquitously expressed lncRNA (*LINC00116*) encodes a highly conserved 56-amino acid mitochondrial transmembrane microprotein, named Mitoregulin (Mtln)^1, 2^. Mitochondria are dubbed the powerhouses of the cell, generating ATP via oxidative phosphorylation (OXPHOS), wherein electrons are shuttled along electron transport complexes (ETC) (cI, cII, cIII, cIV) to O_2_ the final electron acceptor. Apart from OXPHOS, mitochondria also have fundamental roles in Ca^2+^ buffering, lipid and amino acid metabolism, ROS generation and redox signaling, as well as regulated cell death pathways^3^.

The mitochondrion has a distinct double-membrane arrangement, with a porous outer mitochondrial membrane (OMM) and a selectively permeable inner mitochondrial membrane (IMM) that separates the internal matrix compartment from the intermembrane space (IMS). The IMM is highly invaginated to form the cristae tubules, studded with OXPHOS complexes^4, 5^. Mitochondrial membrane lipid composition is distinguished from other cellular membranes by a paucity of sterols and sphingolipids and the presence of the unique tetra-acylated phospholipid, cardiolipin (CL). The major lipids of the OMM are phosphatidylcholine (PC; 54%) and phosphatidylethanolamine (PE; 29%). The IMM requires a high proportion (approximately 50%) of conical shaped lipids (PE and CL), concentrated at negative curvatures, critical to forming and stabilizing cristae shape^5, 6^. Cristae shape in-turn is vital to support efficient OXPHOS operation and is dynamically altered to meet metabolic demand^4, 7^.

Mtln shows a high correlation of co-expression with OXPHOS proteins, particularly accessory proteins necessary for ETC supercomplex assembly^1^. In line with this, loss of Mtln impacts mitochondrial respiration and ROS production^1, 8–11^. In addition, several studies support that Mtln affects various other pathways and phenotypes, including lipid metabolism^2, 8, 12–14^, mitochondrial-ER contacts and ER stress signaling^10^, cancer cell migration/invasion^15^, skeletal muscle performance^2, 9, 13^, myocyte differentiation or regeneration^9, 16^, and normal^17^ and diseased kidney^18^ phenotypes.

Mtln has been found to exist in both the IMM^1, 2, 9^ and OMM^14^, and several diverse Mtln interacting proteins have been reported. Pulldown of overexpressed epitope-tagged Mtln from cell or tissue lysates followed by mass spectrometric analyses uncovered alpha and/or beta subunits of the fatty acid β-oxidation (FAO) trifunctional protein (Hadha and Hadhb)^2, 12^, the alpha subunit of mitochondrial F_1_F_0_ATP synthase (Atp5a)^12^, or cytochrome b5 reductase 3 (Cyb5r3)^8^ as Mtln interactors. Most recently, carnitine palmitoyl transferase 1b (Cpt1b) was co-immunoprecipitated with endogenous Mtln^14^. Additionally, biomolecular fluorescence complementation screening in HTC75 fibrosarcoma cells and HEK293T cells identified cI subunit Ndufa7^15^ and N-acetyltransferase 14 (Nat14)^18^, respectively, as top hits. Despite these findings, a consensus molecular function for Mtln remains elusive. Given that Mtln is a small membrane protein devoid of annotated canonical protein interacting / structural domains yet strongly binds CL^1^, it is plausible that the primary function of Mtln is modulate mitochondrial membranes, potentially assisting in the formation or stabilization of lipid microdomains and cristae structure. In this way, Mtln could secondarily secure various proteins or protein complexes within these domains. This could account for broad, indirect protein interactions and effects of Mtln across several mitochondrial pathways, several of which have strong relevance to various disease conditions.

Mitochondria are vital to supporting the energy-demanding maintenance of normal cardiac function and heath. However, in the context of myocardial ischemia and subsequent reperfusion (I/R), mitochondria contribute to driving pathological events. During the ischemic phase, lack of oxygen shuts down OXPHOS, and cells rely on anaerobic glycolysis, leading to cytosolic acidification and rapid depletion of energy stores. With sustained ischemia, cardiomyocyte cell death will occur in proportion to ischemic time, consequent to energy crisis and inability to sustain ion pumps^19^. Restoration of blood flow, though necessary to salvage cardiac function, introduces Ca^2+^ overload, ROS, and inflammatory conditions that drive additional cardiomyocyte death, collectively referred to as reperfusion injury^20^. Ca^2+^-induced, ROS-facilitated opening of the mitochondrial permeability transition (mPT) pore (mPTP) is widely accepted as an early reperfusion event that precipitates substantial cardiomyocyte death^20–22^. Mitochondria have a relatively large capacity to buffer Ca^2+^, but if Ca^2+^ uptake exceeds a threshold (i.e. maximum retention capacity), opening of the mPTP ensues, allowing unrestricted, bidirectional passage of solutes of up to 1500 Da across the IMM. This dissipates the electrochemical gradient, depletes solutes from the matrix, and allows for H_2_O influx into the protein-rich matrix, promoting matrix swelling and ultimately OMM break. Moreover, mPTP opening destabilizes supercomplexes and compromises ETC function to exacerbate ROS^23^. Various cell death pathways may be invoked; OMM release of pro-apoptotic factors and ROS-mediated events can intersect and contribute to apoptosis or regulated necrotic pathways such as necroptosis and ferroptosis^19, 20^. Although the exact molecular identity of the mPTP is controversial^24^, strong candidates are the adenine nucleotide translocator (Ant) ^25^ and the F_1_F_0_ATP synthase^26^, which may be triggered to reassemble into mPTP. In addition, established facilitators include cyclophilin D, mitochondrial Ca^2+^ and ROS, inorganic phosphate, and free fatty acids ^27^.

Our previous observations show that Mtln overexpression enhances mitochondrial Ca^2+^ retention capacity and resistance to mPT in cells, whereas Mtln-KO mouse myofibers display reduced Ca^2+^ retention capacity^1^. In addition, Mtln gain- and loss-of-function in cells led to decreased and increased mitochondrial ROS levels, respectively. Together, these data support that Mtln could also play a critical role in protecting cardiomyocytes from mPT and I/R injury.

In this current study, we took steps towards elucidating the influence of Mtln on mitochondrial membranes by investigating mitochondria susceptibility to freeze/thaw damage in global Mtln-KO mice and in mice with acute Mtln KO or overexpression (i.e. rescue) compared to controls and by evaluating Mtln’s influence on synthetic IMM-like lipid monolayers. Furthermore, we performed LC-MS lipidomic analyses to identify alterations in lipid profiles of Mtln deficient cardiac tissues and mitochondrial isolates. Lastly, we examined the effect of Mtln deficiency on cardiac outcomes after surgically-induced myocardial I/R injury. Together, our results support a model in which Mtln interacts with mitochondrial membrane lipids to protect CL and reinforce cristae structure, while loss of Mtln enhances mitochondrial vulnerability to membrane insults and I/R injury.

## RESULTS

### Mitochondria from Mtln-KO mice are more susceptible to freeze/thaw-induced membrane damage

In our original publication, we performed blue native gel electrophoresis (BNGE) with mitochondria isolated from mouse heart and observed selective loss of matrix protein signals from Mtln-KO samples compared to wildtype (WT) samples, despite equal amounts of these proteins in total tissue lysates^1^. In those experiments, tissues were collected and stored at -80°C prior to mitochondrial isolation. In follow-up BNGE experiments, we compared mitochondria isolated from fresh versus previously frozen heart tissue. Immunoblot for matrix protein complexes (Ogdh, Acadvl) after transfer from blue native gels shows indistinguishable banding patterns for Mtln-KO and WT mitochondria from fresh tissue; however, if tissues were frozen prior to mitochondrial isolation, Mtln-KO samples showed reduced matrix protein levels compared to WT controls (**Fig. 1A**). This is consistent with our previous observation and supports the notion that enhanced loss of soluble matrix protein complexes from Mtln-KO mitochondria is a consequence of increased freeze damage to mitochondrial membranes.

**Figure 1.**
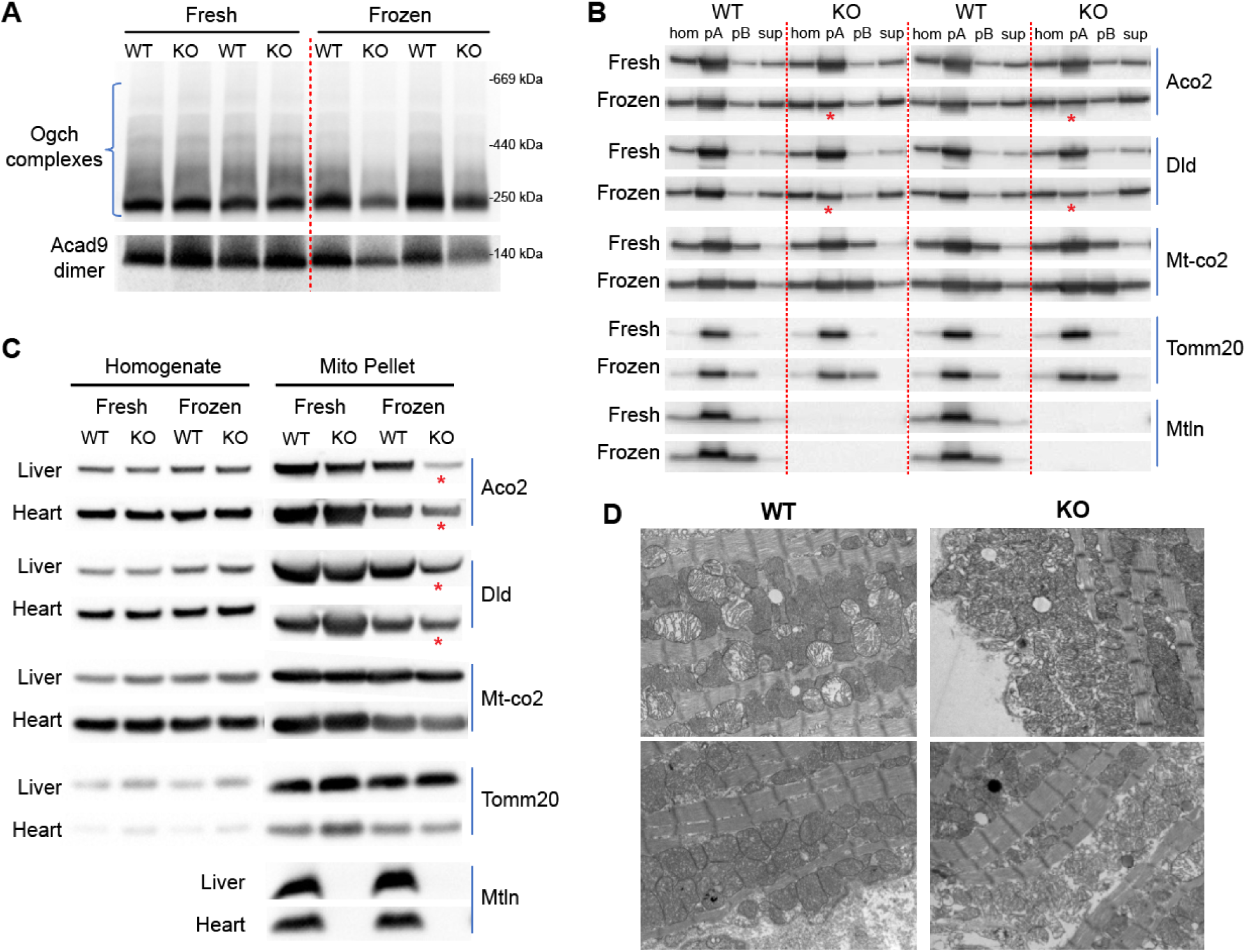
Mitochondria from Mtln-KO compared to WT mice are more susceptible to freeze/thaw-induced membrane damage and matrix protein leak. **A**, Western blot after blue native gel electrophoresis (BNGE) of mitochondria isolated from fresh or frozen/thawed heart tissue from wild-type (WT) and Mtln knock-out (Mtln-KO) mice. N=2 mice each group. **B,** Western blot of samples collected during mitochondria isolation from fresh or frozen/thawed heart tissue from WT and Mtln-KO mice: homogenate (hom), mitochondrial pellet A (pA), mitochondrial pellet B (pB), final supernatant (sup). N=2 mice each group. **C,** Western blot of homogenates and corresponding mitochondrial pellets isolated from fresh or frozen/thawed heart or liver tissue from WT and Mtln-KO mice. N=2 mice each group. **D,** Representative TEM images of heart samples collected after freeze/thaw from WT or Mtln-KO mice. Oxoglutarate dehydrogenase (Ogdh); acyl coenzyme-A dehydrogenase 9 (Acad9); aconitase 2 (Aco2); dihydrolipoamide dehydrogenase (Dld); mitochondrial cytochrome c oxidase subunit 2 (Mt-co2); translocase of outer mitochondrial membrane 20 (Tomm20); Mitoregulin (Mtln). Ogdh, Acad9, Aco2 and Dld are soluble mitochondrial matrix proteins.

We investigated this freeze/thaw phenotype more thoroughly by standard denaturing SDS-PAGE and western blot assay. For this, heart tissues from WT and Mtln-KO mice were minced then divided into “fresh” versus “frozen” (snap frozen/thawed) parts, prior to homogenization. The clarified homogenate (Hom) was centrifuged to generate mitochondrial pellets (pA) by standard differential centrifugation. Overlying supernatants were collected for a second round of centrifugation at higher speed to amass damaged mitochondria in secondary pellets (pB), and the final supernatants overlying pB (Sup) were also collected. Samples were subjected to SDS-PAGE and immunoblot to determine levels of proteins representing different mitochondrial compartments. When isolated from fresh tissue, immunoblot bands are nearly identical for WT and Mtln-KO samples, revealing no significant differences across compartments (**Fig. 1B**). In contrast, when tissues were frozen prior to mitochondrial isolation, exacerbated soluble matrix protein loss (Aco2 and Dld) was evident in pA mitochondria from Mtln-KO samples versus WT controls, and this was accompanied by stronger appearance of these bands in Sup fractions (**Fig. 1B**). In a repeat experiment, we confirmed this phenotype in heart tissues, and further replicated this effect in liver mitochondria isolated from Mtln-KO versus WT mice (**Fig 1C**). Ultrastructural examination of heart tissue by transmission electron microscopy (TEM) after freeze/thaw revealed regions with swollen mitochondria and loss of matrix staining density for both Mtln-WT and KO (**Fig. 1D**). Consistent with our inferences from the western blot data, swollen mitochondria as well as mitochondria with severe membrane discontinuity and fracture were more pronounced in Mtln-KO samples (**Fig. 1D**). Thus, mitochondria lacking Mtln are more susceptible to freeze/thaw-induced membrane damage and resultant leakage of matrix proteins.

Weakened mitochondrial membranes in Mtln-KO mice might arise from loss of a direct Mtln-stabilizing effect or may develop after accumulation of lipid damage and remodeling that occurs over time. In fact, a recent report shows onset of kidney dysfunction and pathology in aged (∼2 years-old) but not young Mtln-KO mice, and those authors propose dysfunction develops after chronic CL oxidation and remodeling^17^. To determine whether our phenotype could develop in a shorter timeframe, we used the CRISPR-Cas9 system to establish acute Mtln-KO in young mice. For this we delivered AAV9 encoding Mtln-targeted guide RNAs (AAV9:gu) or control AAV9:GFP to αMHC-Cas9 transgenic mice, wherein the αMHC promoter drives cardiomyocyte-restricted Cas9 expression^28^. We first validated the effectiveness of this model and achieved a remarkable degree of Mtln elimination at 4 weeks post injection (**Fig. 2A**), with the residual (approximate 5%) protein potentially arising from non-cardiomyocyte cell types, given that Mtln is ubiquitously expressed across cells and tissues. With experimental cohorts, we isolated mitochondria from fresh versus frozen samples collected at 4 weeks post AAV administration, and assayed samples by western blot. Akin to constitutive Mtln-KO samples, acute Mtln-KO also rendered mitochondria more vulnerable to matrix protein leakage after freezing, compared to controls (**Fig 2B**, red asterisks). As expected, this was not observed in liver tissue, which lacks Cas9 expression in these mice (**Fig S1**). Together, these results indicate that the membrane vulnerability in the absence of Mtln is not reliant on long-term accumulated damage and point to a more direct or proximal influence of Mtln on mitochondrial membrane integrity.

**Figure 2.**
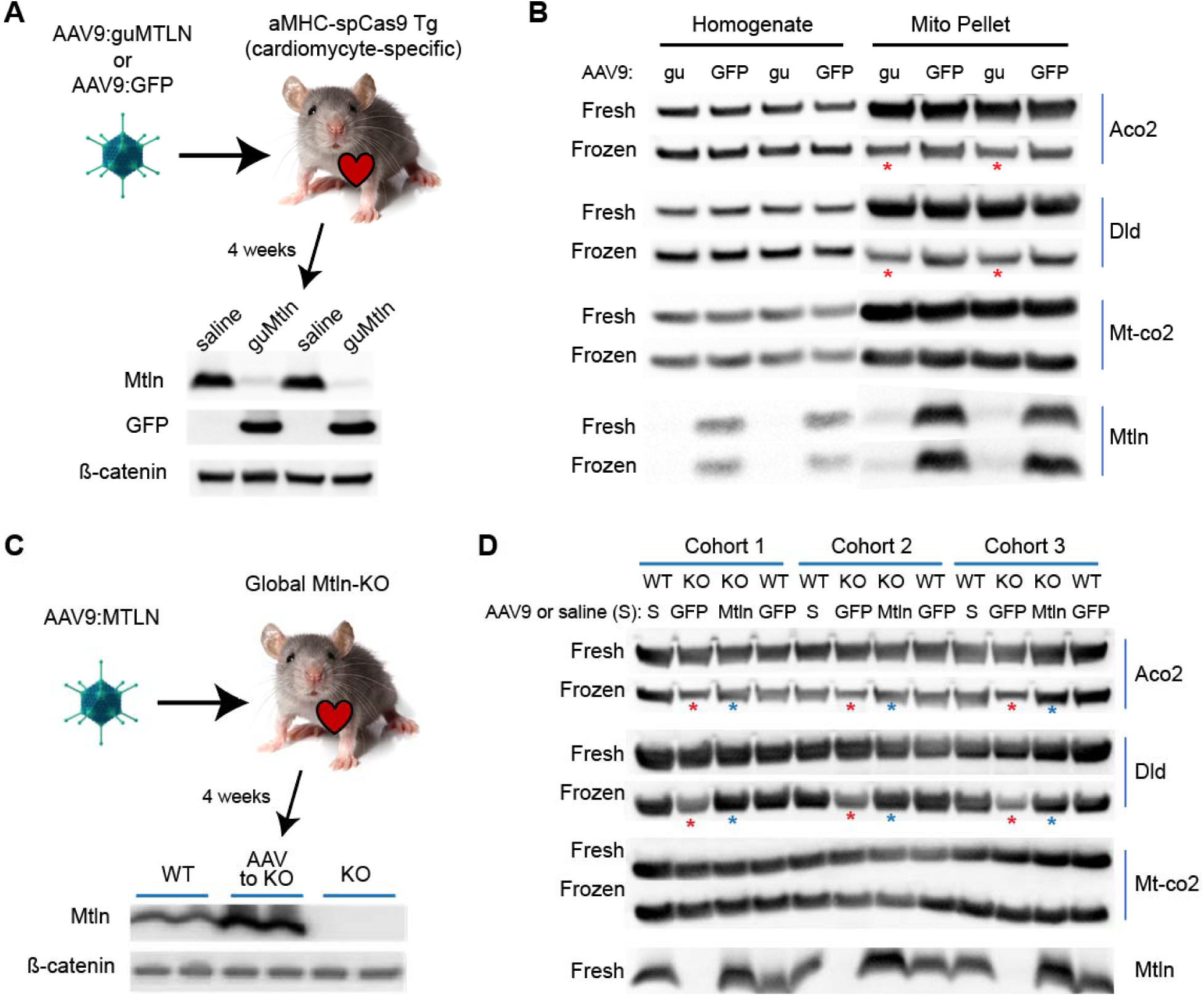
Acute modulation of Mtln expression in mouse heart is sufficient to influence mitochondrial susceptibility to freeze/thaw damage. **A,** Experimental design schematic shown along with western blot comparison of Mtln level in heart tissue collected from aMHC-Cas9 transgenic mice at 4 weeks post intravenous injection of either AAV9:GFP- or AAV9:guMTLN, at 1.0E+10 vg/g mouse weight. N=2 mice each group. **B,** Western blot of homogenates and corresponding mitochondrial pellets isolated from fresh or frozen/thawed heart tissues from AAV9:GFP- and AAV9:guMTLN-injected aMHC-Cas9 transgenic mice. N=2 mice each group. **C,** Experimental design schematic shown along with western blot comparison of Mtln level in heart tissue collected from non-injected WT, non-injected Mtln-KO, or Mtln-KO mice at 4 weeks post intravenous injection of 1.0E+10 vg/g AAV9:Mtln. N=2 mice each group. **D,** Western blot of homogenates and corresponding mitochondrial pellets isolated from fresh or frozen/thawed heart tissue from saline-injected WT mice, AAV9:GFP-injected WT mice, AAV9:GFP-injected Mtln-KO mice, or AAV9:Mtln-injected Mtln-KO mice, with vector dose of 1.0E+10 vg/g. N=3 mice each group.

To further address if this phenotype relates to more immediate Mtln actions versus distal downstream consequences of prolonged Mtln deficiency, we tested if acutely restoring Mtln expression could protect mitochondria from freeze/thaw damage. For this, we used recombinant AAV9 to add back Mtln to Mtln-KO mouse hearts. We first tested the efficacy of this strategy to replenish Mtln and found that AAV9:Mtln restored myocardial Mtln protein to ∼1.5-fold compared to WT levels (**Fig. 2C**). We next assessed whether Mtln add-back would protect against exacerbated freeze/thaw damage. Notably, at 4 weeks post AAV administration, we observed complete rescue of normal phenotype with AAV9:Mtln treatment, replicated in three separate mouse cohorts (**Fig. 2D**). With tissues post-freeze, matrix proteins (Aco2 and Dld) are severely diminished in mitochondria from control (AAV9:GFP-injected) Mtln-KO mice (**Fig. 2D**, red asterisks) but are preserved (to WT level) in mitochondria from Mtln-restored (AAV9:Mtln-injected) KO hearts (**Fig. 2D**, blue asterisks). These data provide further evidence that Mtln protein acutely confers strength to mitochondrial membranes to prevent damage and matrix protein leakage. We postulate that the newly synthesized Mtln may insert into membranes of existing mitochondria to generate membrane alterations that enhance resistance to freeze/thaw-induced membrane damage and likely influence mitochondrial architecture in other important ways.

### Acute loss of Mtln in cells disrupts mitochondrial cristae

To further assess if Mtln acutely influences mitochondrial membranes and ultrastructure, we performed TEM after Mtln knockdown achieved via siRNA transfection into A549 cells, which express high levels of endogenous Mtln. At 48 h post-transfection, Mtln protein levels were reduced by >80% relative to siRNA control (**Fig. 3A**). By TEM, mitochondria were characterized in blinded fashion as either healthy (abundant, aligned cristae tubules), unhealthy (few or disorganized cristae), or intermediate (**Fig. 3B**, representative images). In cells with Mtln silencing, there was a clear and significant shift towards more unhealthy mitochondria with reduced cristae density (**Fig. 3B,C**; p<0.0001). Malformation of cristae within 48 h of Mtln loss in A549 cells boosts the supposition that Mtln provides direct modulatory effects on mitochondrial membranes and ultrastructural integrity.

**Figure 3.**
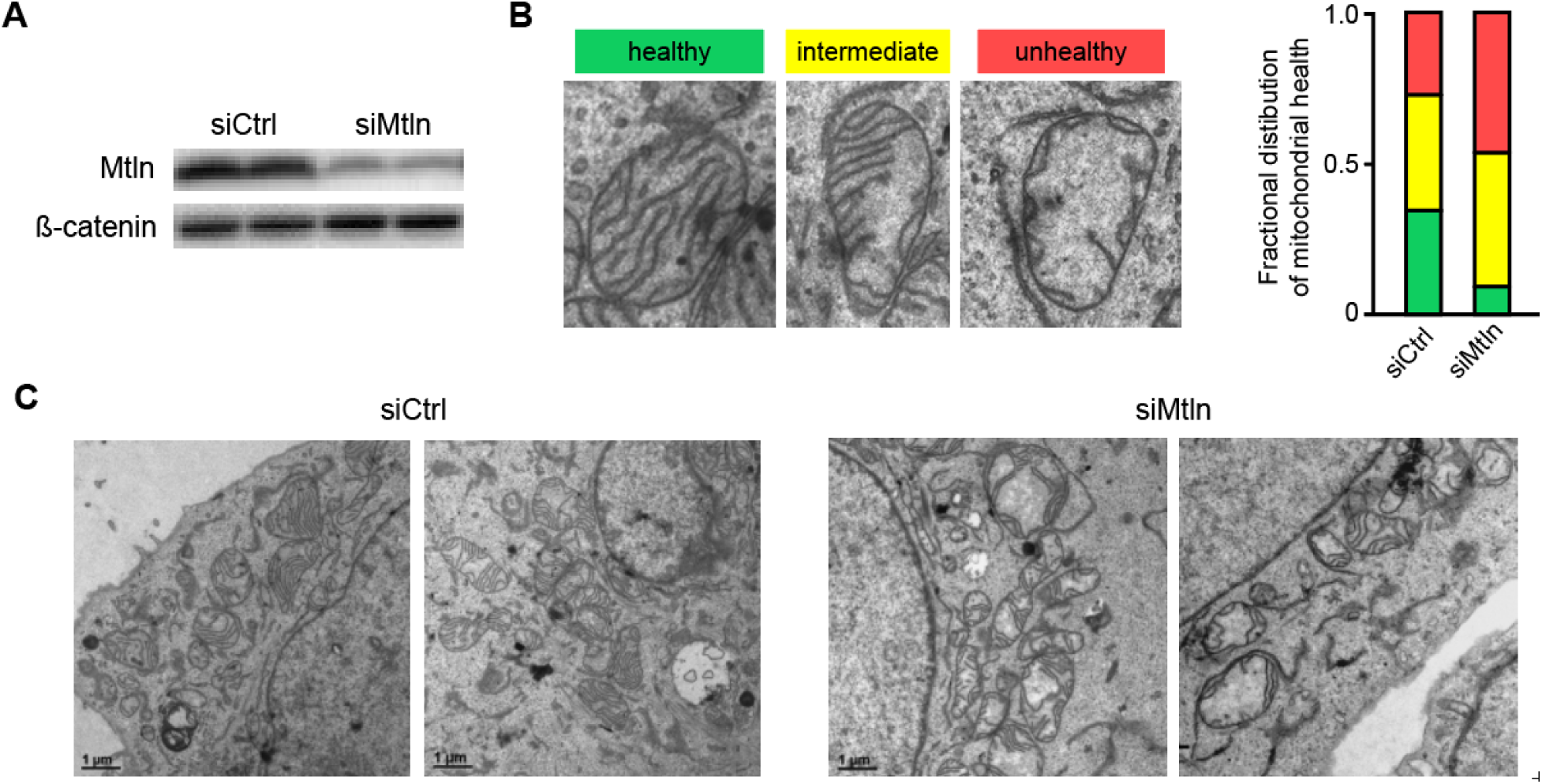
Acute silencing of Mtln in A549 cells disrupts mitochondrial cristae structure. **A,** Western blot for Mtln protein level at 48 h post introduction of negative control (siCtrl) or Mtln-specific (siMTLN) small-interfering RNA oligos. N=2 per group. **B,** Left, schematic with representative TEM images to illustrate cristae scoring system. Right, bar chart showing the distribution of healthy/intermediate/unhealthy mitochondria at 48 h post siCtrl or siMTLN treatment; N=3 transfected cultures per group, with ∼100 total mitochondria scored (from at least 3-4 cell per well) per group. Fisher exact test indicates a significant (p<0.0001) shift toward unhealthy mitochondria in siMtln-treated cells, relative to siCtrl. **C,** Representative TEM images from cell monolayers at 48 h post siCtrl pr siMTLN treatment.

### Mtln peptide directly modulates dynamics of mitochondrial-simulated lipid monolayers

To assess direct influence of Mtln on membrane properties, we applied synthetic Mtln to biomimetic mitochondrial-like monolayers generated in a Langmuir trough. With this instrument, nanomole amounts of defined lipid mixtures are dispersed onto an aqueous surface and a lateral barrier compressed the lipids until they interact to form a monolayer at the liquid-air interface (Langmuir film). As the film is further compressed, molecular packing proceeds and surface pressure rises (surface tension drops), with output profiles dependent on the lipid composition. For each compression run, a lipid mixture (mimicking the IMM) was dispersed onto the surface and Mtln was injected into the buffer subphase to final concentrations of 0.28 μM or 2.8 μM, or no protein was injected. The Langmuir pressure-area isotherms (**Fig. 4A**) illustrate the relationship between surface pressure and the mean area per lipid molecule within the monolayer. The addition of 0.28 μM Mtln shifts the curve slightly to the right, towards a larger “area per lipid molecule”, and this effect is greatly pronounced with 2.8 μM Mtln. Notably, these are significant increases, shown at a physiologically relevant surface pressure of 30 mN/m (**Fig. 4B**). This signifies interaction of Mtln with the monolayer in a manner that creates a less packed overall lipid arrangement. This could be a consequence of Mtln interacting with phospholipid head groups to create lipid conformational changes, and/or due to Mtln inserting more deeply into hydrophobic regions to alter lipid packing within acyl chains.

**Figure 4.**
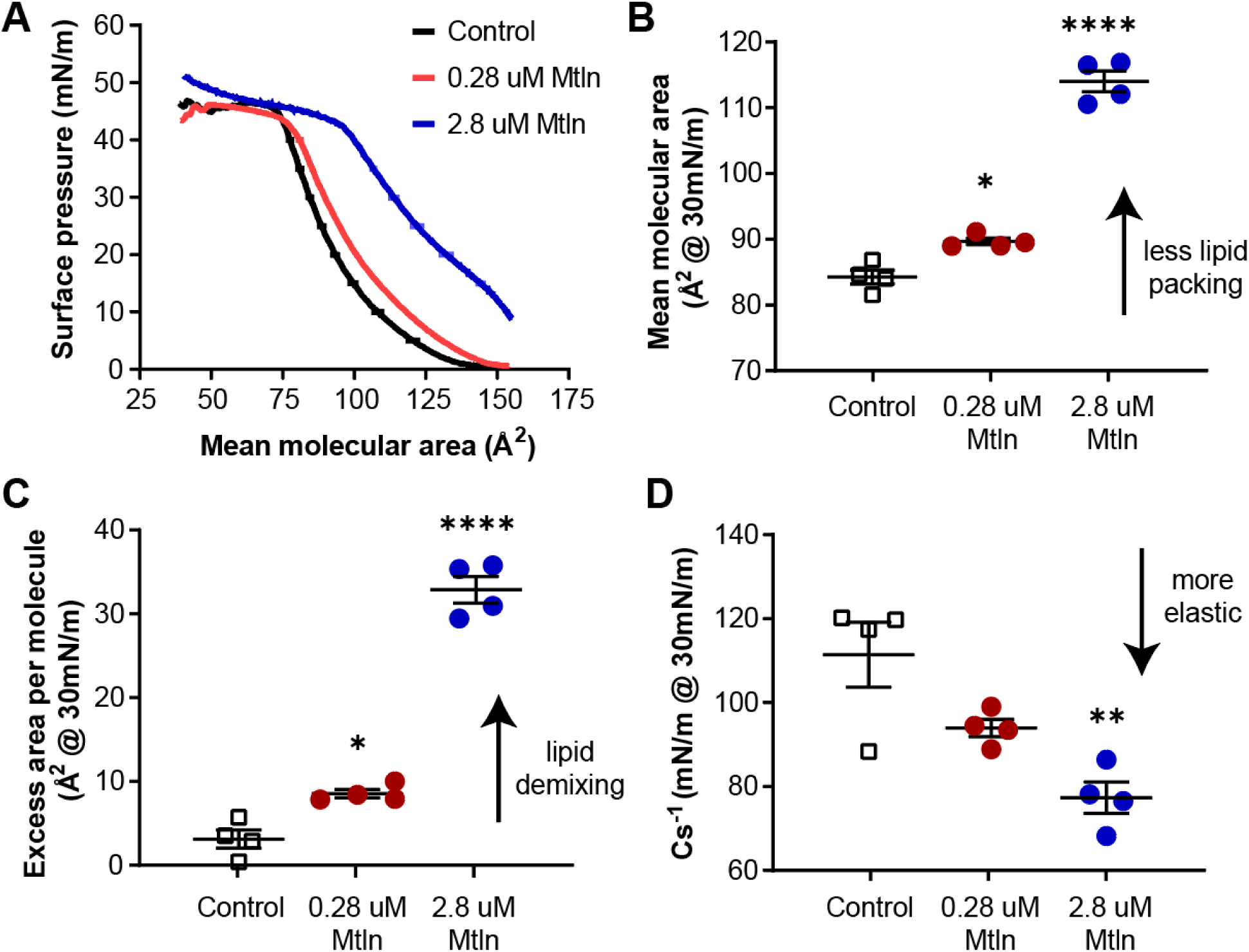
Synthetic Mtln protein interacts with biomimetic mitochondrial-like lipid monolayers and alters membrane properties. **A,** Langmuir trough pressure-area isotherms generated from a mitochondrial biomimetic lipid mixture, with or without the addition of synthetic Mtln to the buffer subphase. N=4 per group. **B,** Plot showing mean molecular area, generated from isotherm raw data at the physiologically relevant surface pressure of 30 mN/m. **C,** Plot showing excess area per molecule (ExA), calculated from the isotherm data at a surface pressure of 30 mN/m. **D,** Plot showing elastic modulus of area compressibility (Cs^-1^), calculated from the isotherm data at a surface pressure of 30 mN/m. Plots show the mean±SEM for N=4 per group. P-values were obtained using one-way ANOVA with Tukey’s multiple comparisons test.

Lipid miscibility can be inferred from the excess area per molecule (ExA). For a given surface pressure, the ExA is the difference between the area per molecule of the lipid mixture (acquired from the pressure-area isotherms) and the ideal area per molecule (expected if lipids behaved independently). Data from mono-lipid isotherms are used to derive weighted-average ideals of the mixture. A negative ExA indicates attractive forces, while a positive ExA suggests inter-lipid repulsive forces and resistance to lipid mixing. At the surface pressure of 30 mN/m, we found that the IMM-like lipid film without Mtln registers a slightly positive ExA (**Fig. 4C**), indicating close to ideal lipid miscibility with a small introduction of repulsive forces. With increasing addition of Mtln, the ExA is significantly more positive in a dose-responsive manner (**Fig. 4C**). This outcome suggests that adsorption of Mtln to the monolayer (and/or into the monolayer via hydrophobic interactions) causes lipid de-mixing. With a polybasic C-terminal tail, Mtln is expected to interact electrostatically with headgroups of anionic lipids (especially CL), similar to cationic SS-31 peptide interaction with model membranes^29^. As was previously observed with cytochrome C (CytC)^30^, Mtln interaction could trigger segregation of lipids species, to explain the rise in ExA.

Lastly, the isotherm raw data was used to further evaluate Mtln’s possible influence on membrane dynamics by calculating the elastic modulus of area compressibility (Cs^-1^), which is the inverse of compressibility. At physiologic 30 mN/m surface pressure, Cs^-1^ decreases with addition of Mtln, indicating increased compressibility or elasticity of the lipid monolayer (**Fig. 4D**). Together these Langmuir film analyses show that Mtln interacts with and/or adsorbs into IMM-like monolayers and imparts changes in lipid spacing, mixing, and elasticity, all of which are fundamental to IMM folding, cristae formation and membrane lipid dynamics, and would plausibly affect membrane response to freeze/thaw or other stresses.

### Mtln-KO mouse hearts have elevated triglycerides and decreased docosahexaenoic acid

To assess Mtln’s influence on lipid profiles, cardiac tissue collected from male Mtln-KO and WT mice (∼21-month-old) was subjected to high performance liquid chromatography tandem mass spectrometry (LC-MS). Use of positive and negative ionization modes for MS provided broad lipidome coverage, identifying 436 unique lipid species. Mtln-KO samples show a clear pattern of increased triglycerides (TG), particularly those containing saturated or lowly desaturated acyl chains of 14-20 carbons in length (**Fig. 5A** and **Table S1**). Among the 42 significantly altered lipids identified (p≤0.05, Mtln-KO versus WT), 35 are elevated TG species. By contrast, lipid species containing 22:6(n-3) (docosahexaenoic acid, DHA) acyl chains, including 22:6-containing TG, are mostly reduced in Mtln-KO samples (**Fig. 5A**).

**Figure 5.**
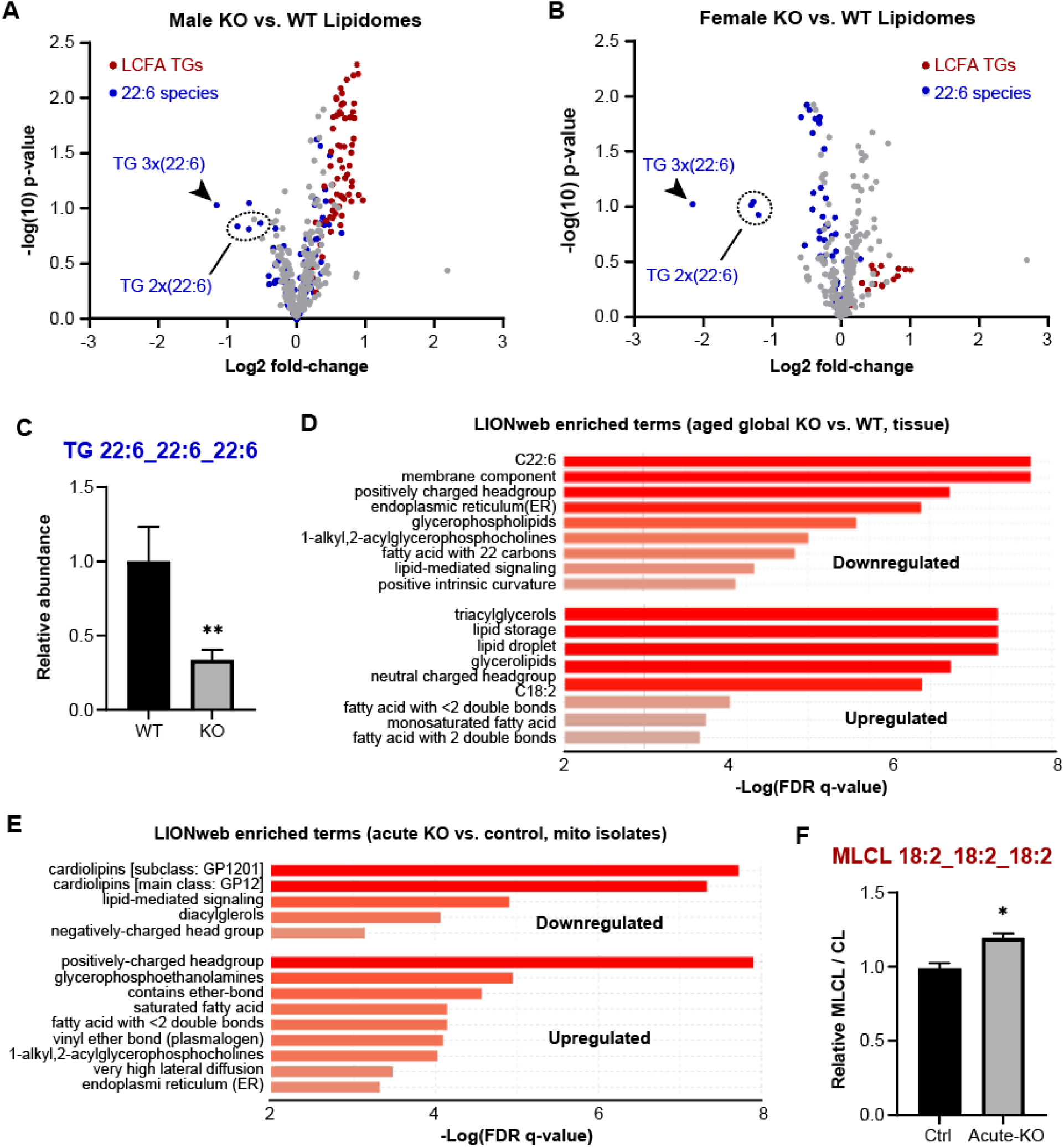
Aged Mtln-KO mouse hearts have elevated triglycerides and decreased docosahexaenoic acid while acute Mtln loss in young hearts triggers cardiolipin (CL) reduction / remodeling. **A-D,** Lipid species from heart apex of aged (21 to 23 month-old) WT and Mtln-KO mice were quantified by high performance liquid chromatography tandem mass spectrometry (LC-MS). **A,** Volcano plot display of univariate analysis for males (N=5 each group). **B,** Volcano plot display of univariate analysis for females (N=5 each group). **C,** Bar graph showing relative abundance of TG 22:6_22:6_22:6 in WT compared to Mtln-KO mice, mean±SEM (N=10 per group, 5 for each gender). **D,** Lipid ontology enrichment (LiON) analysis for Mtln-KO relative to WT lipidomes (N=10 per group, 5 for each gender). **E and F,** Lipid species from mitochondria isolated from aMHC-Cas9 transgenic mice at 4-weeks post AAV9:GFP or AAV:guMtln injection (7 weeks of age at sample harvest), were quantified by LC-MS; N=6 male mice each group. **E.** LiON analysis for AAV9:guMtln relative to AAV9:GFP lipidomes. **F,** Bar graph showing monolysylcardiolipin/cardiolipin (MLCL/CL) ratios (mean±SEM) for AAV9:GFP and AAV9:guMtln samples. P-values were obtained using unpaired two-tailed student’s t-test.

Subsequent lipidomic profiling was done on heart samples collected from aged WT and Mtln-KO female mice, once again revealing an enrichment of phospholipids with saturated or low desaturation level (including TGs, albeit to a lesser degree than males) in Mtln-KO samples, as well as a significant reduction in highly desaturated lipid species (acyl chains with desaturation ≥5, **Fig. 5B** and **Table S2**), which is more pronounced than in males. Notably, among all lipids detected, TG containing three (3x) and two (2x) 22:6 acyl chains shows the strongest reduction (by magnitude) in both female and male Mtln-KO samples (**Fig. 5A,B**). Gender-combined analysis shows that TG 22:6_22:6_22:6 is reduced by ∼65% in Mtln-KO samples (**Fig. 5C**, p=0.01). These general findings (decreases in 22:6 species and increase in other TG) were corroborated by both lipid ontology enrichment analysis (LIONweb, **Fig. 5D**) and unsupervised, multivariate *partial least squares discriminant analysis* (PLS-DA, data not shown). Together, these lipidomic studies done on bulk cardiac tissue samples reinforce that Mtln loss causes a shift away from FAO^1, 2, 12^ towards increased TG lipid storage^12^ and highlight an accompanying selective broad reduction in lipids with highest desaturation levels.

### Acute cardiac-specific Mtln-KO causes cardiolipin turnover

To address specifically how loss of Mtln acutely influences mitochondrial lipids, we performed lipidomic profiling on crude mitochondrial pellets isolated from fresh cardiac tissue from conditional Mtln-KO mice (αMHC-Cas9 transgenic mice injected with AAV9:guMtln) or control mice (αMHC-Cas9 transgenic mice injected with AAV9:GFP), collected 4 weeks post-AAV) (model shown in **Fig. 2C**). Among 544 individual lipid species quantified, few show significant differences between groups (**Table S3**). Targeted examination of DHA (22:6)-containing lipids revealed weakly trending differences (p<0.2), encompassing 15 unique lipids, all of which are notably down-regulated, consistent with our findings in bulk cardiac tissues. Interestingly, lipid ontology enrichment analysis (LIONweb, **Fig. 5E**) indicated that acute Mtln-KO samples showed broad reductions in CL species, as well as accumulation of lipids having “head group with positive charge”, glycero-ethanolamines, and plasmalogens (vinyl ether phospholipids), the latter of which could represent an attempt to preserve IMM curvature, as these lipid classes encompass several alternative conical lipid species (like CL). Moreover, the *sn*-1 vinyl ether linkages of plasmalogens are highly reactive to ROS and serve as scavengers to prevent peroxidation of other lipids^31, 32^. This is notable since prior observations indicate that mitochondrial ROS rises after Mtln knockdown in cells^1, 10^, and the plasmalogen spike after acute Mtln reduction could serve as an early compensatory antioxidant defense. Acute Mtln-KO mitochondria showed additional signs of ROS-induced membrane damage, with lipidomics data indicating a weak trend towards increases in monolysocardiolipin (MLCL 18:2_18:2_18:2, p<0.2), and notably, follow-up auxiliary analyses revealed that MLCL/CL ratios are significantly increased by ∼20% in acute Mtln-KO samples (**Fig. 5F**, p=0.015, representing the “top hit”, most significant p-value in this lipidomics dataset). Such an increase in MLCL/CL ratio represents a classic hallmark of CL damage/turnover, as occurs in Barth syndrome^33^. Along with this key finding, the overall mitochondrial-targeted lipidomics data provide complementary insights into the multimodal means by which Mtln may broadly influence lipid composition and integrity to maintain membrane stability through freeze/thaw, and this may translate further to relevant pathological insults that cause mitochondrial membrane damage and remodeling.

### Mtln-KO mice exhibit worse myocardial injury in response to ischemia-reperfusion

As an initial step towards evaluating a possible protective role for Mtln in disease, we chose to investigate a mouse model of myocardial ischemia-reperfusion (I/R), given that Mtln is cardiac-enriched and that I/R is known to elicit significant and pathologic mitochondrial membrane disruption. Ischemia and/or reperfusion events can trigger mitochondrial permeability transition to promote cell death pathways^20^. Moreover, mitochondrial swelling, loss of matrix staining density, and cristae fragmentation are detected in mouse heart after I/R^34^. We first assessed cardiac function of Mtln-KO mice compared to WT in the absence of stress and found that baseline cardiac function (determined by echocardiography) is normal, for both young (∼15-20 weeks-old) and middle-aged (>1-year of age) mice **(Fig. 6A, Tables S4 and S5**). To examine if absence of Mtln would affect outcomes of I/R, mice were subjected to coronary artery ligation, blocking blood flow to the lower left ventricular wall for 1 h, after which the suture was removed to restore blood flow. Given that we hypothesized that Mtln-KO mice would exhibit exacerbated I/R injury, we performed surgeries using polypropylene suture, which in our hands (versus nylon), limits the severity of ischemia and tissue damage, providing a better window of opportunity to observe worse outcomes in mice that may be more susceptible to I/R injury. We performed surgeries on young male mice (Mtln-KO and WT littermates, ∼18 weeks of age), and at 3 days post-surgery, heart structure and function were measured by echocardiography (**Fig. 6B**; summary data in **Table S6**). Among 13 WT mice, only 5 (38%) showed discernible infarcts in echocardiographic images, whereas 7 of 11 Mtln-KO mice (64%) had obvious infarcts, which on average were nearly twice the size compared to WT infarcts (**Fig. 6C**, p<0.05). Overall, Mtln-KO mice showed significantly worse outcomes, evidenced by increased dilation (larger end-diastolic and end-systolic volumes) and reduced left ventricular ejection fraction, relative to WT mice (**Fig. 6C**). These data provide initial evidence supporting a protective role for Mtln in I/R injury, though the complex and likely interrelated underlying mechanisms (involving combined influences on membrane stability, mitochondrial Ca^2+^ and ROS, and mPTP) will need to be resolved with extensive follow-up investigations.

**Figure 6.**
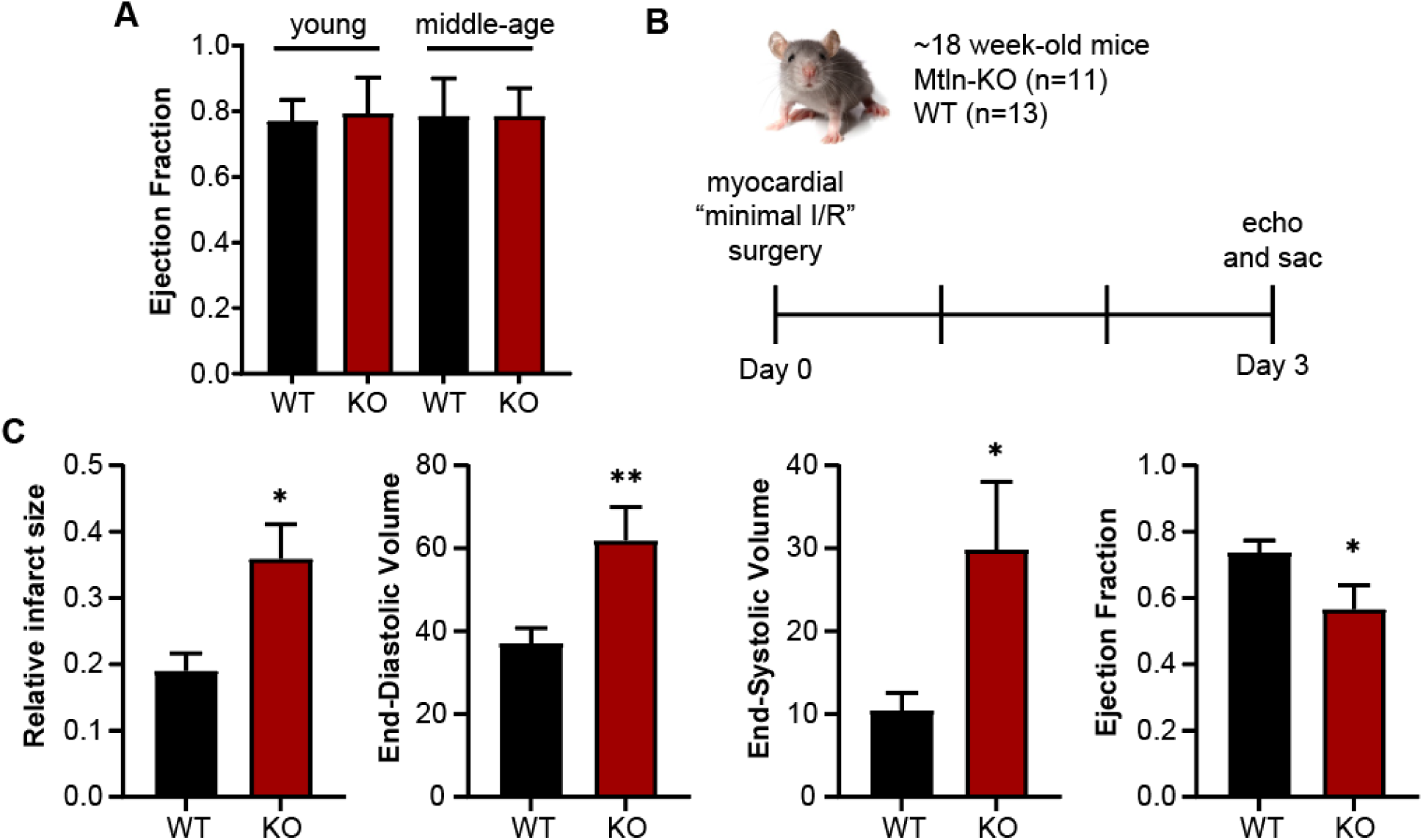
Mtln-KO mice exhibit worse myocardial injury in response to ischemia-reperfusion (I/R). **A,** Echocardiographic measurement of baseline ejection fractions mice at two age-groups, young (15-20 weeks old; N=6 WT, N=6 Mtln-KO), and middle-age (1-1.3 y old N=11 WT, N=13 Mtln-KO). **B,** Workflow diagram of myocardial I/R experimental approach. **C,** Bar graphs of significantly changed echocardiographic measurements collected at 3 days post reperfusion; N=13 WT, N=11 Mtln-KO). P-values were obtained using unpaired two-tailed student’s t-test.

## DISCUSSION

This study was spurred by an intriguing difference that we serendipitously observed between WT and Mtln-KO mitochondria after isolating mitochondria from frozen cardiac tissue. We discovered striking exacerbation of matrix protein leakage from Mtln-KO compared to WT mitochondria and verified this phenotype for mitochondria isolated from liver as well as heart. TEM assessment of frozen/thawed cardiac tissue revealed regions with swollen or damaged mitochondria, which were in greater abundance and appeared more highly fractured in Mtln-KO tissues, raising the interesting questions of 1) what is happening during freeze/thaw, 2) how does lack of Mtln worsen the outcome, and 3) what other genes and/or physiologic contexts (e.g. aging, obesity, mitochondrial disease) influence the magnitude of mitochondrial freeze/thaw damage?

It is well established that subjecting cells or tissues to freeze/thaw induces mitochondrial membrane damage, which translates to worse respiratory function. While newer protocols have been developed to enable measures of mitochondrial respiration in frozen tissues^35^, which offers significant conveniences, our work highlights the need to consider whether possible differences in the extent of freeze/thaw damage across experimental groups could confound these methods. This is likely to be influenced by a breadth of biological conditions and processes. Without cryoprotectant compounds, substantial damage to membranes is expected, as membranes are sensitive to osmotic changes and dehydration effects of ice crystal formation^36, 37^. Dramatic mitochondrial swelling after freeze/thaw is documented in mammalian cells^38^ and in cold-susceptible insect larvae^39^. Mitochondrial swelling occurs after loss of the IMM barrier integrity and subsequent osmotic influx of water into the protein-rich matrix. Theory asserts that with temperature drop/dehydration upon freezing, and with temperature rise/rehydration upon thaw, membrane lipids experience unregulated and asynchronous liquid-gel phase transitions and separations^40^. This creates regions with increased tendency to form non-lamellar structures, such as reversed hexagonal phase and inverted micelles, that can ablate barrier function^40, 41^, essentially creating pores. Transient formation of such non-lamellar structures is known to occur normally under physiologic conditions to promote membrane fusion or fission^42, 43^. Lipids prone to non-lamellar arrangements are more abundant in organelles with tubular structure such as the ER and IMM^44^. CL is one such lipid having well-known propensity to form transient reverse hexagonal phases^43, 45, 46^.

Mtln strongly binds CL^1^, and our Langmuir analyses demonstrate that Mtln adsorbs onto (and/or into) mitochondrial-like lipid monolayers and alters monolayer properties. Mtln reduces lipid density and enhances elasticity. We propose that one physiologic role of Mtln *in vivo* is to interact with CL in microdomains that support membrane folding and cristae structure. In the context of freeze/thaw, Mtln intercalation in CL rich regions could antagonize lipid phase transitions and CL propensity to form reverse hexagonal structures that create pore-like openings. Additionally, increased membrane elasticity imparted by Mtln may provide resistance to swelling-induced fracture, to allow eventual volume recovery. Other events, such as rise in Ca^2+^ concentration, due to water withdrawal into ice crystals and/or leakage from damaged ER, may contribute essential steps to excessive formation of non-lamellar structures. In fact, when added to CL preparations^45^ or to CL-containing vesicles^47, 48^ or nanodiscs^49^, Ca^2+^ promotes inverted hexagonal structures and vesicle leak. Importantly, CL levels after acute Mtln-KO, though slightly below WT samples, would still be of sufficient abundance to generate non-lamellar formations. Further investigation is necessary to better define the molecular mechanisms as to how Mtln concerts with CL, possibly to restrain transitions to reverse-hexagonal lipid arrangements and provide resistance to membrane break upon freeze/thaw.

We cannot exclude the possibility that Mtln has indirect effects on mitochondrial lipids to impact vulnerability to temperature stress. Lipid composition influences tendency for phase separations. For example, a study in bacteria demonstrated that raising the content of saturated membrane lipids (tight packing, low fluidity) precipitated striking de-mixing and phase separation of the inner membrane lipids into low and high fluidity regions^50^. Our lipidomic results show broad lipid alterations upon Mtln loss, and one interesting finding is the drop in lipid species containing 22:6 in Mtln-KO. The 22:6 fatty acid, docosahexanoic acid (DHA), is an omega-3 fatty acid and is one of the most highly desaturated fatty acids. Phospholipids with 22:6 chains would enhance bilayer fluidity and perhaps lessen phase separations that contribute to freeze/thaw damage in highly unsaturated membranes. However, the drop in 22:6 species was more drastic at the tissue-level versus mitochondrial-level lipidome. This could mean that loss of 22:6 from additional cellular compartments contributes indirectly to IMM damage, or that decline in 22:6 is not a key determinant of exacerbated mitochondrial membrane damage in Mtln-KO. Either way, the mechanism for how Mtln influences DHA-containing lipids remains unclear. Notably, two independent reports demonstrate that Mtln-KO cell lines exhibit elevated DHA or DHA-containing lipid species^8, 14^, hinting at indirect effects of Mtln loss on fatty acid metabolism that are context-dependent (*in vitro* versus *in vivo*).

An important translationally relevant finding in our current study is that Mtln supports cardiac resilience to I/R. The detrimental effects of I/R were heightened in the absence of Mtln; infarct zone was enlarged along with increasing cardiac dilation and impaired performance. Widely accepted as a prominent contributor to reperfusion injury is mitochondrial permeability transition (mPT), which occurs at the onset of reperfusion during a critical period of high Ca^2+^ and mitochondrial ROS surge^20^. Above a threshold, matrix Ca^2+^ triggers formation and opening of the mPT pore (mPTP), a non-selective, high conductance (up to 1500 Da) IMM pore of incompletely defined molecular identity^24, 27^. Pore opening collapses the mitochondrial membrane potential, causing energy crisis, water influx into the matrix, mitochondrial swelling, OMM rupture, and trigger of cell death pathways^20^. Recent studies support transition of either the Ant an/or the F_1_F_0_ATP synthase (or a component of the F_1_F_0_ATP synthase), from its normal structure/function into pathologic mPTP formation^24, 27^. Pore opening can be regulated by cyclophilin D, and threshold to opening is substantially lowered by ROS^24, 27^. Given that Mtln acutely enhances mitochondrial Ca^2+^ retention capacity^1^ (a measure of resistance to permeability transition), it is possible that Mtln antagonizes cardiac mPTP opening after reperfusion, to protect a proportion of at-risk cardiomyocytes from dying. We speculate that Mtln binding to CL may also play a direct part in resistance to mPT. Though the field is still controversial regarding molecular events and protein components of mPTP^24, 27^, both Ant- and F_1_F_0_ATP synthase-central models incorporate possible sensor roles for CL. CL is an integral component of these protein complexes^51, 52^ and CL shifts conformation when exposed to Ca^2+^ or when oxidized, and in this way may behave as a sensor to elicit mPTP configuration^53, 54^. In line with this, Ca^2+^ alters CL packing in lipid bilayers and renders CL more susceptible to oxidation^55^. CL loss occurs after cardiac I/R^34, 56^, indicative of ROS induced oxidation and subsequent turnover^56^. By virtue of binding to CL, Mtln feasibly shields CL from Ca^2+^ effects and/or limits CL acyl chain exposure to I/R-induced ROS.

Recent 3D ultrastructural examination of cardiac mitochondria documents mitochondrial swelling, loss of matrix staining density, and fragmentation of cristae connectivity early after I/R^34^. Intriguingly, cristae connectivity was significantly preserved by application of the therapeutic mitochondria-targeting cationic tetrapeptide, SS-31 (elamipretide) during reperfusion^34^. SS-31 targets IMM CL, to stabilize CL-integral protein complexes^57, 58^ and demonstrates broad mitochondrial supportive effects under various pathologic conditions^59–61^. Notably, SS-31 mediates protection against hypoxia-driven lipid peroxidation of CL^58^. SS-31 benefits are reminiscent of Mtln, such that Mtln, which similarly contains “basic plus aromatic amino acid” segments, may represent an endogenous SS-31-like counterpart, binding to CL to provide direct support against various stresses.

Our working model tying together freeze/thaw and I/R protective effects (**Fig. 7**), predicts that Mtln partners with CL to stabilize CL conformation in a manner that i) limits propensity to form membrane-damaging, non-lamellar structures upon temperature stress; and ii) shields against oxidation and cleavage upon I/R stress. We found evidence for broad CL turnover in heart mitochondria after acute removal of Mtln, consistent with a role for Mtln in stabilizing CL in a manner that restricts acyl chain access. Concordant with this finding, CL decline and increased MLCL/CL ratio was recently described in skeletal muscle mitochondria from Mtln global KO mice^13^. Unlike their observation in the soleus, our findings were less robust, and we found no significant drop in CL or rise in MLCL in heart tissue from our global Mtln-KO mice. We suspect that with sustained absence of Mtln, the heart adapts by introducing altered lipid species to provide some CL stability and/or by altering CL remodeling pathways. Once formed, CL-protein complexes, and CL-protein supercomplexes are particularly stable in cardiomyocytes^62, 63^; a mild change in CL turnover might go undetected in the non-stressed heart that has adapted to Mtln loss. With I/R stress, the rise in mitochondrial ROS is predicted to result in oxidation of CL, while ROS and Ca^2+^ directly stimulate iPLA2γ^64, 65^ to cleave CL/oxidized CL, generating MLCL. We predict that these events are exacerbated in the absence of Mtln (**Fig. 7**). Interestingly, when complexed with CL or MLCL, CytC acquires (per)oxidase activity^66, 67^; CL and MLCL are themselves targets of oxidation and serve to propagate peroxidation of additional polyunsaturated membrane lipids for extended membrane damage^67^. Thus, we surmise that events downstream of I/R-induced CL turnover are intensified in the absence of Mtln, and include mPTP assembly, and MLCL-CytC association to perpetuate lipid peroxidation (**Fig. 7**). At this juncture, our finding that Mtln deficiency worsens I/R injury is of great interest and will steer our future efforts towards deeper interrogation of our model to better understand the molecular function of Mtln in I/R injury pathways.

**Figure 7.**
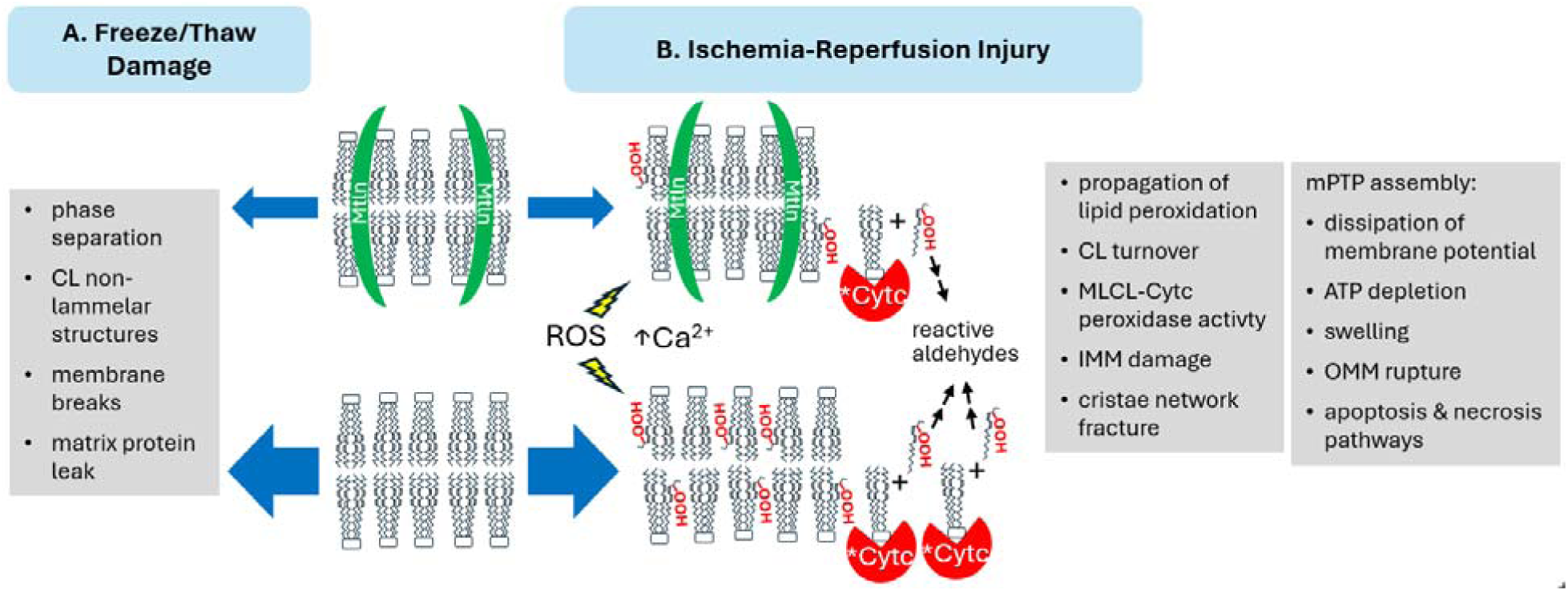
Model: Mitoregulin association with CL shields against mitochondrial membrane damage under stress. Working model shows Mtln interacting directly with CL to: **A)** reduce tendency for CL to self-associate into damaging non-lamellar structures upon freeze/thaw and **B)** shield CL from peroxidation and degradation during cardiac ischemia-reperfusion and thus limit predicted outcomes such as propagation of extensive membrane peroxidation, mPTP formation and trigger of cell death pathways.

Our lipidomic analyses indicate a general increase in TG content at the tissue and mitochondrial level. Other investigators have reported increased TG^8^ or lipid droplets^12^ in Mtln KO cells, indicative of a diversion of lipids away from mitochondrial FAO and towards TG storage. This could reflect a direct involvement of Mtln in either lipolysis^12^ or FAO^2, 12, 14^, the latter of which is consistent with reported Mtln interactions with FAO related proteins^2, 12, 14^. Another thought is that OMM and IMM structural deformations in the absence of Mtln is poorly supportive of FAO, perhaps due to ineffective intersection of FA uptake and FAO enzymatic complexes at OMM-IMM boundaries. An alternative explanation is that diminished FAO is an indirect outcome as part of metabolic rewiring to compensate for impaired mitochondrial respiration, downstream of abnormal CL. Interestingly, metabolic re-wiring accounts for blunted FAO observed in cells deficient in tafazzin^68^, the CL-remodeling transacylase mutated in Barth syndrome. These possibilities await more in-depth evaluations.

We expect that Mtln influence on IMM lipid dynamics, particularly involving CL, would have broad effects on metabolism and cell functions related to mitochondria, and could explain the multifaceted outcomes of Mtln loss across studies. It is also possible that Mtln additionally functions in a key manner within defined protein complexes, for example with trifunctional proteins and Cpt1b for FAO^2, 12^ or assisting Cyb5r3 oxidoreduction reactions^8^, or with Nat-14 to mediate inflammation-triggered nuclear signaling^18^. Mtln may perform diverse functions and localize to more than one mitochondrial subregion or protein complex, in a cell-type or tissue-specific manner and/or depending on metabolic demands.

Collectively, our study advances knowledge of Mtln influence on mitochondrial membranes, highlighting the importance of Mtln in direct interaction with IMM lipids, and a connection to CL turnover. Moreover, our findings reveal a role for Mtln in cardiac stress responses, specifically in blunting I/R injury. While the translational significance of this remains unclear, further studies could be warranted to better understand how environmental or physiologic influences affect Mtln expression in the heart, as well as assess whether Mtln overexpression in mouse heart is sufficient to confer protection against myocardial I/R injury. Our continued molecular and functional investigations will strive to provide enlightenment on the operational mechanisms of this intriguing microprotein. Resolving Mtln’s mode-of-action in countering stress and promoting mitochondrial health has potential to inspire discovery of new targets for I/R therapy and likely other mitochondrial-related pathologies.

## ACKNOWLEDGEMENTS

The transgenic aMHC-cas9 mouse line was graciously provided by Eric Olson (UTSW). This work utilized the Hitachi S-4800 Scanning Electron Microscope in the University of Iowa Central Microscopy Research Facilities that was purchased with funding from the NIH SIG S10 RR022498; the content is solely the responsibility of the authors and does not necessarily represent the official views of the NIH. We acknowledge the Metabolomics Core at the University of Utah and thank James Cox and John Alan Maschek for assistance and help with lipidomic analyses. for their assistance in (service provided). Echocardiography was performed with the help of the University of Iowa Cardiovascular Phenotyping Core directed by Dr. Robert Weiss with technical support from Kathy Zimmerman, Alyssa Bosko, and William Kutschke. This work was supported by the University of Iowa Carver College of Medicine (Distinguished Scholars Program to R.L.B), NIH NHLBI (HL148796, HL144717, and HL150557 to R.L.B.), NIH NIGMS (predoctoral fellowship T32 GM067795 to N.H.W), and the American Heart Association (23PRE1011277 to N.H.W.).

## AUTHOR CONTRIBUTIONS

R.L.B. conceived and designed the project, supervised the research, and analyzed and interpreted the data. C.S.S, X.Z., N.H.W., and E.R.P. designed and executed experiments, curated and analyzed data, and participated in data interpretation. S.R.S. assisted with experimental design and data interpretation. C.S.S. and R.L.B. wrote the manuscript with contributions from X.Z., N.H.W., E.R.P. and S.R.S.

## MATERIALS AND METHODS

### Mouse studies

All animal studies and procedures were approved by the Institutional Animal Care and Use Committees (IACUC) at the University of Iowa. Mice were housed in a controlled temperature environment on a 12 h light/dark cycle, with food and water provided ad libitum, unless otherwise noted. Mtln knock-out (Mtln-KO) mice were generated as previously described^1^ at the University of Iowa Genome Editing Facility using CRISPR/Cas9 technology, deleting the mouse short open reading frame (1500011K16Rik gene). Mtln-KO mice were back-crossed 9 or more generations to C57BL6/J or C67BL6/NJ (Jackson Laboratory stock numbers #000664 and #005304, respectively). The Myh6-Cas9-2A-TdTomato transgenic mouse (aMHC-Cas9), with Cas9 driven by the cardiomyocyte alpha myosin heavy chain (aMHC) gene (Myh6) promoter was kindly provided by Eric Olson^28^, back-crossed for >9 generations to C57BL/6J, and heterozygous aMHC-Cas9 mice were used for acute knock-out experiments.

### Cell culture and transfection

Human A549 lung carcinoma cells (ATCC, CCL-185) were maintained in Gibco DMEM/F-12 (Thermofisher 11320033) with 100 U/ml penicillin, 0.1 mg/ml streptomycin and 10% FBS in a cell culture incubator in standard conditions (humidified, 37°C with 5% CO2). For Mtln silencing, a dicer-ready 27-mer RNA oligo pair targeting the human Mtln open reading frame, 5’rArCrUrGrCrArGrUrUrGrUrCrCrGrUrGrCrUrArGrUrArGCC3’ and 5’rGrGrCrUrArCrUrArGrCrArCrGrGrArCrArArCrUrGrCrArGrUrGrU3’, was generated from Integrated DNA Technology (IDT, Coralville IA) using their dsiRNA design tool. Cells were transfected with 50 nM Mtln-directed (siMTLN) or negative control (siCtrl) (IDT cat# 51-01-14-03) dsiRNA oligos, using Lipofectamine Ltx (Invitrogen) according to product instructions. At 48 h post transfection, cell monolayers were processed for either western blot or transmission electron microcopy (TEM). For western blot, media was removed, and lysis buffer (50 mM TRIS-HCL, pH 7.5 with 1% LDS) with 1x Halt protease inhibitors (Invitrogen) added to monolayers. Lysed samples were collected to tubes on ice, and probe sonicated (Fisher Scientific, Sonic Dismembrator Model D100) for two rounds of 5 s each at intensity level 3. Protein concentrations of lysates were determined by BCA assay (Pierce BCA Protein Assay Kit, Thermofisher Ref# 23225), equilibrated, and western blot performed as described below. For TEM, media was removed, TEM fixative (2.5% glutaraldehyde in 0.1M sodium cacodylate buffer, pH 7.2) was added to monolayers and the plates were wrapped in foil and stored at 4°C till processing for TEM as described below.

### Mitochondria isolation

Reagents and tubes were ice-cold, tissue processing was performed on ice, and centrifugations were done at 4°C. Tissues were dissected from mice under deep anesthesia, or immediately after euthanasia. Tissue samples (heart, with atria removed, or liver lobe) were finely minced in a small volume of isolation buffer (in mM, 250 sucrose, 5 Tris-HCL pH 7.4, 2 EGTA, 5 MgCl_2_, 1 ATP, and 0.5% fatty acid-free BSA, with final adjustment to pH 7.4 before use). Minced tissue was transferred to isolation buffer and pelleted by 1 min centrifugation at 800 × G. After discarding the supernatant, the minced tissue was resuspended in isolation buffer with 0.2% BSA (3-5 ml), transferred to an 8 mL Potter Elvehjem tissue grinder with PTFE Pestle (Kimble 886000–022) and homogenized manually for 6-9 passes. After pelleting debris at 800 × G for 10 min the clarified homogenate (Hom) was transferred to a fresh tube and centrifuged at 8000 × G for 10 min to obtain the mitochondrial pellet. For “fresh vs frozen” experiments, the minced tissue was divided to two tubes in isolation buffer and centrifuged at 1 min at 800 × G to pellet the minced tissue, and the supernatant discarded. For the “fresh” sample, the minced tissue was resuspended and homogenized for mitochondrial isolation as described above. For the “frozen” sample, the minced tissue was snap-frozen by tube immersion in LN_2_, thawed by brief warming at 37°C, then further processed in identical fashion as the fresh sample. For “fresh vs frozen” experiments, the BSA content in the isolation buffer during homogenization was lowered to 0.01%, to enable accurate protein determination of Hom samples. For “fresh vs frozen” experiments, a small aliquot (90 ul) of clarified homogenate (Hom) was collected (before the 8000 x G spin step), and for some experiments the supernatant over the mitochondrial pellet (pellet A, pA) was transferred to a fresh tube and spun at 20,000 x G for 20 minutes to generate a second (damaged mitochondria) pellet (pB), and a final supernatant (Sup). Hom, mitochondrial pellets, and Sup samples were snap frozen and stored at -80 until processing for gel electrophoresis, western blotting, or lipidomics.

### Blue native gel electrophoresis (BNGE) and transfer to PVDF

BNGE procedures were derived from published methods^69, 70^. Mitochondrial pellets were resuspended in aminocaproic acid buffer (ACAB, 50 mM bis-tris, pH 7.0 with 0.75 M 6-aminocaproic acid) with freshly added protease inhibitors (Roche). After resuspension, a small aliquot was removed, lysed in 1% Triton X-100, and used for protein concentration determination by BCA. Resuspended mitochondrial samples were then brought to equal concentrations in ACAB buffer, and digitonin (high purity, Millipore) was added from a 10% stock to give a 4 g:g ratio of (digitonin:mitochondrial protein) and samples were solubilized on ice for 20 min, then centrifuged for 20 min at 20,000 × G at 4°C. Pelleted debris was discarded and supernatants (solubilized mitochondrial proteins or protein complexes) were transferred to fresh tubes and coomasie brilliant blue G (CBBG, Sigma B0770) was added from a 5% stock to give a 4 g:g (digitonin:CBBG) ratio. Glycerol was added to a final 5%, and equal volumes containing equal total protein were loaded into wells of precast NativePAGE Novex® 3%–12% Bis-Tris 10-well Gels (Invitrogen). High molecular weight (HMW) native markers (GE Healthcare) reconstituted in 100 μL ACAB plus protease inhibitors and 5 ul 5% CBBG, were loaded at 10–15 μL per lane. BNGE was run in an Invitrogen Novex mini-Cell tank in a 4°C cold room. Cathode buffer (15 mM bis-tris, 50 mM tricine,) and anode buffer (50 mM bis-tris) were pH 7.0 at 4°C. CBBG was added to the cathode buffer to a final 0.02% (dark blue) or 0.0005% (light blue). Electrophoresis was initiated at constant 150V with dark blue cathode buffer, and after approximately 50 min the dark blue was replaced with light blue cathode buffer and electrophoresis resumed at 250V for a maximum of 3 h total running time (until the dye front reached the end of the gel). Gels were transferred to 0.2 μm PVDF membrane using the Invitrogen Mini Blot Module and NuPAGE transfer buffer at 4°C at 30V overnight in a BioRad mini-blot wet transfer system. After transfer, membranes were fixed for 10 min in 8% acetic acid, rinsed in water and air-dried. HMW marker lanes were cut off, and the remaining membrane was re-wet in methanol and washed in several changes of methanol to remove CBBG, equilibrated in water for 2 minutes, then blocking and immunodetection steps were followed as described below for standard western blot. The HMW marker lanes were re-wet briefly in methanol until bands were clearly visible, then transferred to water, and re-aligned with the blot during imaging.

### Standard denaturing PAGE and western blot

For “fresh vs frozen” mitochondrial isolation experiments, Hom, mitochondrial pellets, and Sup samples (as defined in Mitochondrial isolation) were thawed on ice. Protease inhibitors (Roche) and triton x-100 were added (to final 1x and 1% concentrations, respectively) to Hom and Sup samples. Mitochondrial pellets were resuspended in lysis buffer (50 mM TRIS-HCL, pH 7.5 with 1% LDS) with added protease inhibitors. Within each experiment, Hom protein concentrations were determined by BCA assay and Hom concentrations were equilibrated. Sup and mitochondrial lysate volumes were adjusted to the same relative degree as their corresponding Hom. After addition of NuPage sample buffer (denaturing and reducing), samples were heated for 10 minutes at 50°C and equal volumes were resolved on 4-12% bis-tris gradient NuPage gels with MES buffer or MOPS running buffer (Invitrogen) and transferred to 0.2 μm PVDF membrane (Millipore) using NuPAGE transfer buffer (Invitrogen) with added 10% ethanol and 0.01% SDS. After transfer, membranes were blocked by 1 h room temperature incubation in tris-buffered saline with 0.1% tween (TBST) containing 5% milk (dissolved dry milk powder), then incubated overnight with primary antibodies at 4°C on an orbital shaker. Primary antibodies were diluted in either TBST with 2.5% milk or TBST with 0.05% BSA. Primary antibodies were: anti-Mtln (1/2000–1/3000, custom Proteintech specific Mtln c-terminal amino acids 38-56)^1^, anti-OGDH (1/2000, Abclonal A6391), anti-ACAD9 (1/4000, Aviva OAAN02440), anti-VDAC1 (1:2000, Abclonal), anti-β-catenin (1:2000; Sigma-Aldrich, PLA0230), anti-DLD (ABlclonal A13296). Blots were washed in TBST and incubated with HRP-conjugated secondary antibody from goat (1:20000 – 1:60000; Jackson ImmunoResearch, 115–035–146 and 111– 035–144) in blocking buffer, washed in TBST and developed using ECL-prime (GE Health-care) or SuperSignal Femto (ThermoScientific) and chemiluminescent images captured on a VersaDoc Imaging system with Quantity One software (Bio-Rad) or iBright imaging system.

### Transmission electron microscopy (TEM)

For TEM of cardiac samples after freeze/thaw, Mtln-KO and WT mice were fasted for 4 h, anesthetized with ketamine/xylazine, and the heart apex was snipped off and transferred to a metal block on ice with a few drops of mitochondrial isolation buffer (without BSA), and cut into <1mm pieces. The pieces were transferred to a tube and flash frozen in LN_2_, thawed quickly by hand, and immediately transferred to TEM fixative (2.5% glutaraldehyde in 0.1M sodium cacodylate buffer, pH 7.2) and stored at 4°C protected from light. For TEM preparation, the fixed heart samples, or fixed A549 monolayers (prepared described in “Cell culture and transfection”, were washed with 0.1M sodium cacodylate buffer and stained with 4% osmium tetroxide and 2.5% uranyl acetate. Samples were washed, dehydrated in graded ethanol washes (50% to 100%), and washed with 100% propylene oxide prior to embedding in EPON resin. Following ultramicrotomy, samples were post-stained with 5% uranyl acetate and Reynold’s lead citrate and examined on a Hitachi HT7800 transmission electron microscope. Digital images were collected with an AMT NanoSprint15 camera (5056×2960 pixel sCMOS sensor).

### AAV vectors

AAV vectors were prepared at the UIOWA Viral Vector Core (VVC). AAV2/9CMVeGFP (AAV9:GFP) was purchased as an in-stock vector (VVC-U of Iowa-238). For custom-ordered AAV2/9CMV-NC116-IRES-GFP (AAV9:NC116), the human NC116 protein coding region was PCR amplified from genomic DNA using primers with added restriction enzyme sites and subsequently inserted into the VVC shuttle plasmid G0692 pFBAAVCMVmcswtIRESeGFPBgHpA using standard cloning techniques. For custom-ordered AAV2/9mU6-sgNC116.2-3-CMVeGFP (AAV9:guMtln), synthetic DNA sequence encoding a validated Mtln-targeted RNA guide pair ^1^, purchased as a DNA g-block (IDT) was inserted into the VVC shuttle vector G0760 pFBAAVmU6mcsCMVeGFPSV40pA using standard cloning techniques. Shuttle plasmids carrying the transgenes of interest were provided to the VVC for triple transfection (3XT) or baculovirus (BAC) methods to generate AAV particles carrying transgenes flanked by AAV2-type inverted terminal repeats (ITRs) and packaged into AAV serotype 9 capsids. Vector particles were purified by iodixanol gradient followed by ion exchange using MustangQ Acrodisc membranes. Vector titers (vector genomes per ml, vg/mL) were determined by digital droplet PCR.

### AAV injection

Mice (3–4 weeks old) were brought to a surgical plane of anesthesia under 1.5-2% isoflurane and 1 liter/minute O_2_ inhalation (Harvard Apparatus Advanced Safety Ventilator). Hair was removed from the surgical site and analgesic (Flunixin or Meloxicam) was subcutaneously injected. A small incision was made through the skin to expose the jugular vein. Under view of a dissecting microscope, AAV particles or vehicle control were injected into the jugular vein using a 30G needle, delivering 1.0E+10 vg per g mouse weight. The incision was closed with 6-0 prolene suture.

### Langmuir Trough Assays

Pressure-area isotherms were generated on a Mini Langmuir-Blodgett Trough (KSV NIMA, Biolin Scientific, Paramus, NJ) using a Wilhelmy plate as previously described^30, 71^. Briefly, a lipid mixture representing inner mitochondrial lipid composition was generated by co-dissolving lipids (40 mol% 18:0–22:6 PC, 33 mol% 16:0–20:4 PE, 20 mol% (18:2)_4_CL, 5 mol% DOPS, 2 mol% cholesterol) in chloroform (10 μg/μl). Lipid monolayers were constructed by spotting 9.0 nmol lipid on a subphase of 10 mM sodium phosphate buffer (pH 7.4). Synthetic human Mtln was synthesized by Biomatik USA, LLC with 95.58% purity, confirmed by the manufacturer with HPLC and mass spectrometry. For experiments where protein was added to the subphase, synthetic human Mtln protein (Genscript), dissolved at 2 mg/ml in water, was immediately injected underneath the lipid monolayer at final concentrations of 0.28 mM or 2.8 mM. Excess chloroform was allowed to evaporate for 10 min prior to monolayer compression. Pressure-area isotherms were generated at a compression rate of 3.0 mm/min and were analyzed at physiologically relevant surface pressures (*i.e.* 30 mN/m) as previously described^30, 71^.

The ideal mean molecular area was calculated at a constant surface pressure (π) by: *A*_ideal_ = *X*_1_(*A*_1_)_π_ + *X*_2_(*A*_2_)_π_ + …*X_n_*(*A_n_*)_π_ (Eq. 1), where *X_n_* represents the mol fraction of each individual component and *A_n_* represents the mean molecular area of each component. The excess area/molecule (*A_ex_*), a measurement of lipid miscibility, was calculated at a given surface pressure (π): *A*_ex_ = (*A*_12…*n*_)_π_ − (*X*_1_*A*_1_+*X*_2_*A*_2_+…*X_n_A_n_*)_π_ (Eq. 2), where *X_n_* represents the mol fraction of each individual component and *A_n_* represents the mean molecular area of each component. (*A*_12…*n*_) represents the mean molecular area of the mixed monolayer of interest at a given surface pressure (π). A positive excess area/molecule indicates repulsive forces, and a negative excess area/molecule indicates attractive forces. Surface pressure-area isotherms were also used to calculate the surface elasticity modulus (Cs^−1^): Cs^−1^=(−A)(drr/dA)rr (Eq. 3), where *A* is the mean molecular area of the lipid mixture of interest at the indicated surface pressure (π).

### LC-MS lipidomics

Mtln-KO and WT mice were fasted for 4-6 h, and while under deep ketamine/xylazine anesthesia, the heart apex was snipped off and immediately snap-frozen by LN_2_ immersion. Apex samples collected from male mice (6 WT and 6 Mtln-KO, 21-22 months old) and female mice (6 WT and 6 Mtln-KO, 21-22 months old) were crushed under LN_2_, weighed, and stored at -80 °C. Mitochondria were isolated from hearts of 6 AAV9:GFP injected and 6 AAV9:guMtln injected aMHC-Cas9 transgenic mice at approximately 7 weeks of age (4 weeks post AAV injection) and mitochondrial pellets stored at -80. Protein content of mitochondrial pellets was determined by BCA assay of distinct, paired pellets (collected from the same homogenate), after detergent solubilization. Apex samples and mitochondrial pellets were shipped on dry ice to the University of Utah Metabolomics Facility for lipid extraction and mass spectrometry.

#### Lipid extraction

Lipid extraction, based on Matyash *et al.*^72^, was performed as follows. All solutions were pre-chilled on ice. Tissues or mitochondrial pellets were transferred to labeled bead-mill tubes (1.4 mm, MoBio Cat# 13113-50) where lipids were extracted in a solution of 250 µL PBS, 225 µL MeOH containing internal standards, and 750 µL MTBE (methyl tert-butyl ether). Internal standards were Avanti SPLASH LipidoMix (Lot#12) at 10 µL per sample and Cambridge Isotope laboratories NSK-B and NSK-B-G1 (deuterated carnitines) at 10 µL per sample. The samples were homogenized in one 30 s cycle using the Omni Bead Ruptor followed by a rest on ice for 1 h. An addition of 188 µL PBS was made to induce phase separation. After centrifugation at 16,000 g for 5 minutes at 4 °C, the upper phases were collected and evaporated to dryness under a gentle nitrogen stream at room temperature. Lipid samples were reconstituted in 500 µL IPA (isopropyl alcohol) and transferred to an LC-MS vial with insert (Agilent 5182-0554 and 5183-2086) for analysis. Concurrently, a process blank sample and pooled quality control (QC) sample was prepared by taking equal volumes (∼50 µL) from each sample after final resuspension.

#### Mass Spectrometry

Lipid extracts were separated on a Waters Acquity UPLC CSH C18 1.7 µm 2.1 x 100 mm column maintained at 65 °C connected to an Agilent HiP 1290 Sampler, Agilent 1290 Infinity pump, equipped with an Agilent 1290 Flex Cube and Agilent 6545 Accurate Mass Q-TOF dual AJS-ESI mass spectrometer. For positive mode, the source gas temperature was set to 250 °C, with a gas flow of 12 L/min and a nebulizer pressure of 35psig. VCap voltage was set at 3500 V, fragmentor at 110 V, skimmer at 65 V and octopole RF peak at 750 V. For negative mode, the source gas temperature was set to 250 °C, with a drying gas flow of 12 L/min and a nebulizer pressure of 30 psig. VCap voltage was set at 3000 V, fragmentor at 100 V, skimmer at 65 V and octopole RF peak at 750 V. Samples were analyzed in a randomized order in both positive and negative ionization modes in separate experiments acquired with the scan range m/z 100 – 1700. Mobile phase A consisted of ACN:H_2_O (60:40 v/v) in 10 mM ammonium formate and 0.1% formic acid, and mobile phase B consisted of IPA:ACN:H_2_O (90:9:1 v/v) in 10 mM ammonium formate and 0.1% formic acid. The chromatography gradient for both positive and negative modes started at 15% mobile phase B then increased to 30% B over 2.4 min, then increased to 48% B from 2.4 – 3.0 min, then increased to 82% B from 3 – 13.2 min, then increased to 99% B from 13.2 – 13.8 min where it was held until 16.7 min and then returned to the initial condition and equilibrated for 5 min. Flow was 0.4 mL/min throughout, injection volume was1 µL for positive and 5 µL for negative mode. Tandem mass spectrometry was conducted using the same LC gradient at collision energy of 25 V.

Data QC samples (n=8) and blanks (n=4) were injected throughout the sample queue to ensure the reliability of acquired lipidomic data. Results from LC-MS experiments were collected using an Agilent Mass Hunter (MH) Workstation and analyzed using the software packages MH Qual, MH Quant, and Lipid Annotator (Agilent Technologies, Inc.). The data table exported from MHQuant was evaluated using Excel where initial lipid targets were parsed based on the following criteria. Only lipids with relative standard deviations (RSD) less than 30% in QC samples were used for data analysis. Additionally, only lipids with background AUC counts in process blanks that were less than 30% of QC were used for data analysis. The parsed excel data was normalized to total lipid signal. For lipid classification visit http://www.lipidmaps.org/data/classification/LM_classification_exp.php.

#### Statistical analysis and pathway mapping

Data for each sample were normalized to sample weight to determine lipid concentrations (pmol per mg tissue). Log2-based fold-change of group means and p-values (using unpaired two-trailed t-test) were determined comparing KO versus WT samples. Gender-combined analyses were done by setting the WT group means for males and female independently to a value of 1 prior to integrating the data. Lipidomes were subjected to unsupervised, multivariate *partial least squares discriminant analysis* (PLS-DA), to reveal lipid features with significant contribution to group separation. Lipid Ontology (LION) enrichment analysis (LION/web; using 2-log local statistics and two-tailed KS-test settings)^73^ were done to identify upregulated and downregulated enriched LION terms for Mtln-KO relative to WT tissues or acute KO (AAV9:guMtln) relative to control (AAV9:GFP) mitochondria.

### Mouse model of minimal myocardial ischemia/reperfusion (I/R) injury

14- to 21- week-old male mice were anesthetized via 1.5% to 2% isoflurane with 1-1.5 liter/minute O_2_ inhalation (Harvard Apparatus Advanced Safety Ventilator), directed through tracheal intubation. The ventilator parameters were adjusted to the body weight of the mouse, to maintain a surgical plane of anesthesia. The third intercostal space over the left chest of the mouse was exposed. Myocardial ischemia-reperfusion injury was induced by ligation of the left anterior descending artery (LAD) with 7-0 prolene suture for 60 minutes, followed by releasing the ligation for reperfusion. Sham-operated mice were treated with the same surgery without tying the left anterior descending coronary artery. Electrocardiography was performed 3 days after reperfusion to evaluate the successful generation of the I/R model. Mouse hearts were then excised for histological and molecular analyses.

### Echocardiography

Echocardiography was performed on conscious mice, with mild sedation (Midazolam 5 mg/kg, SC), using a Vevo2100 imaging system (VisualSonics, Toronto, Canada). Mice were restrained in the left lateral recumbent position in the operator’s hand. Two-dimensional, M-mode, and Doppler imaging were performed in both long- and short-axis views to acquire standard measurements of left ventricular structure and cardiac function according to the endocardial and epicardial area protocol.

### Statistical analyses

Statistical analyses were performed using Prism GraphPad and Microsoft Excel software. GraphPad was used to generate plots and to perform two-tailed t tests or one-way ANOVAs with Tukey’s post hoc to obtain the reported p values. The specific statistical tests for each experiment are indicated throughout the manuscript; for each, all data met the test assumptions and groups showed similar variance. Results were considered to be significant if p values were ≤ 0.05. Statistical significance is directly stated or indicated by convention (* p<0.05, ** p<0.01, *** p<0.001, **** p<0.0001, ns = not significant).

## Supplementary

**Figure S1.**
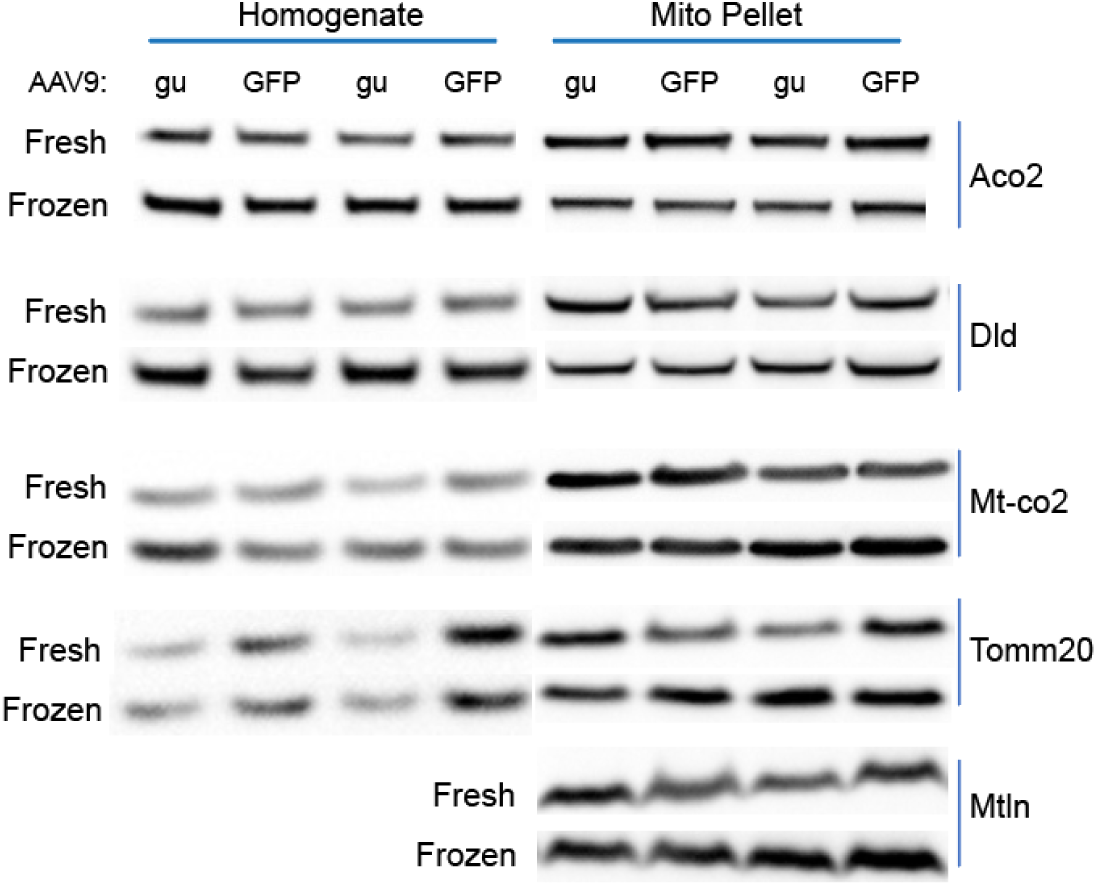
Liver mitochondria are not affected by cardiomyocyte-targeted Mtln knock-out. Western blot of homogenates and corresponding mitochondrial pellets isolated from fresh or frozen/thawed liver tissue from AAV9:GFP- and AAV9:guMTLN-injected aMHC-Cas9 transgenic mice. N=2 mice each group (same mice as in Fig. 2B).

**Table S1.** Summary of LC-MS lipidomic analyses done on mouse hearts collected from aged male WT or Mtln-KO mice. See attached spreadsheet.

**Table S2.** Summary of LC-MS lipidomic analyses done on mouse hearts collected from aged female WT or Mtln-KO mice. See attached spreadsheet.

**Table S3.** Summary of LC-MS lipidomic analyses done on mouse heart mitochondrial isolates collected from young mice with acute Mtln-KO or controls. See attached spreadsheet.

**Table S4.** Summary of echocardiographic measures in young wildtype (WT) and Mtln-KO mice at baseline.

|  | WT |  | Mtl<sup>n</sup>-KO |  |  |
| --- | --- | --- | --- | --- | --- |
|  | Mean | ±SEM | Mean | ±SEM | p-value |
| **Heart rate (bpm)** | 656 | ±24 | 680 | ±15 | 0.41 |
| **ENDO-area (d)** | 7.26 | ±0.50 | 6.13 | ±0.20 | 0.06 |
| **ENDO-area (s)** | 1.79 | ±0.18 | 1.68 | ±0.12 | 0.60 |
| **EPI-area (d)** | 18.36 | ±0.81 | 16.05 | ±0.66 | 0.05* |
| **End-diastolic volume** | 43.85 | ±3.36 | 37.16 | ±1.10 | 0.09 |
| **End-systolic volume** | 8.89 | ±0.86 | 8.20 | ±0.66 | 0.54 |
| **Mass (mg)** | 85.96 | ±3.00 | 77.20 | ±4.41 | 0.13 |
| **Vol/mass** | 0.51 | ±0.03 | 0.49 | ±0.02 | 0.59 |
| **Ejection Fraction** | 0.80 | ±0.01 | 0.78 | ±0.01 | 0.31 |

Young mice (15-20 weeks old; N=6 WT, N=6 Mtl<sup>n</sup>-KO).
Asterisk denotes significant findings comparing the two groups (WT vs Mtl<sup>n</sup>-KO).
ENDO/EPI-areas refer to endocardial/epicardial areas during diastole (d) or systole (s).

**Table S5.** Summary of echocardiographic measures in middle-aged wildtype (WT) and Mtln-KO mice at baseline.

|  | WT |  | Mtl<sup>n</sup>-KO |  |  |
| --- | --- | --- | --- | --- | --- |
|  | Mean | ±SEM | Mean | ±SEM | p-value |
| **Heart rate (bpm)** | 553 | ±35 | 593 | ±43 | 0.48 |
| **ENDO-area (d)** | 8.74 | ±0.59 | 8.13 | ±0.81 | 0.54 |
| **ENDO-area (s)** | 2.59 | ±0.31 | 2.10 | ±0.69 | 0.50 |
| **EPI-area (d)** | 21.20 | ±1.09 | 20.84 | ±1.17 | 0.82 |
| **End-diastolic volume** | 51.88 | ±3.48 | 46.00 | ±4.92 | 0.33 |
| **End-systolic volume** | 12.43 | ±1.70 | 10.39 | ±3.16 | 0.56 |
| **Mass (mg)** | 98.60 | ±4.71 | 96.30 | ±4.62 | 0.73 |
| **Vol/mass** | 0.52 | ±0.03 | 0.48 | ±0.04 | 0.33 |
| **Ejection Fraction** | 0.79 | ±0.02 | 0.79 | ±0.03 | 1.00 |

Middle-aged mice (1-1.3 y old; N=11 WT, N=13 Mtl<sup>n</sup>-KO).
ENDO/EPI-areas refer to endocardial/epicardial areas during diastole (d) or systole (s).

**Table S6.** Summary of echocardiographic measures in wildtype (WT) and Mtln-KO mice collected at 3 days post-I/R.

|  | WT |  | Mtl<sup>n</sup>-KO |  |  |
| --- | --- | --- | --- | --- | --- |
|  | Mean | ±SEM | Mean | ±SEM | p-value |
| **Heart rate (bpm)** | 685 | ±7.8 | 682 | ±16.7 | 0.32 |
| **ENDO-area (d)** | 6.49 | ±0.48 | 10.15 | ±0.98 | 0.0008*** |
| **ENDO-area (s)** | 2.14 | ±0.41 | 5.66 | ±1.24 | 0.01** |
| **EPI-area (d)** | 16.65 | ±0.81 | 20.30 | ±1.07 | 0.004** |
| **End-diastolic volume** | 37.10 | ±3.67 | 57.60 | ±7.97 | 0.007** |
| **End-systolic volume** | 10.45 | ±2.08 | 29.45 | ±8.17 | 0.02* |
| **Mass (mg)** | 75.60 | ±5.00 | 76.04 | ±3.53 | 0.55 |
| **Vol/mass** | 0.48 | ±0.02 | 0.75 | ±0.08 | 0.0006*** |
| **Ejection Fraction** | 0.74 | ±0.04 | 0.51 | ±0.07 | 0.03* |

Young mice (14-21 weeks old; N=13 WT, N=11 Mtl<sup>n</sup>-KO).
Asterisks denote significant findings comparing the two groups (WT vs Mtl<sup>n</sup>-KO).
ENDO/EPI-areas refer to endocardial/epicardial areas during diastole (d) or systole (s).

## REFERENCES

1. Stein, C.S. et al. Mitoregulin: A lncRNA-Encoded Microprotein that Supports Mitochondrial Supercomplexes and Respiratory Efficiency. Cell Rep 23, 3710–3720.e3718 (2018).

2. Makarewich, C.A. et al. MOXI Is a Mitochondrial Micropeptide That Enhances Fatty Acid β-Oxidation. Cell Rep 23, 3701–3709 (2018).

3. Nunnari, J. & Suomalainen, A. Mitochondria: in sickness and in health. Cell 148, 1145–1159 (2012).

4. Cogliati, S., Enriquez, J.A. & Scorrano, L. Mitochondrial Cristae: Where Beauty Meets Functionality. Trends Biochem Sci 41, 261–273 (2016).

5. Ikon, N. & Ryan, R.O. Cardiolipin and mitochondrial cristae organization. Biochim Biophys Acta Biomembr 1859, 1156–1163 (2017).

6. Horvath, S.E. & Daum, G. Lipids of mitochondria. Prog Lipid Res 52, 590–614 (2013).

7. Hu, C. et al. OPA1 and MICOS Regulate mitochondrial crista dynamics and formation. Cell Death Dis 11, 940 (2020).

8. Chugunova, A. et al. LINC00116 codes for a mitochondrial peptide linking respiration and lipid metabolism. Proc Natl Acad Sci U S A 116, 4940–4945 (2019).

9. Lin, Y.F. et al. A novel mitochondrial micropeptide MPM enhances mitochondrial respiratory activity and promotes myogenic differentiation. Cell Death Dis 10, 528 (2019).

10. Choi, M. & Kang, K.W. Mitoregulin controls mitochondrial function and stress-adaptation response during early phase of endoplasmic reticulum stress in breast cancer cells. Biochim Biophys Acta Mol Basis Dis 1869, 166570 (2023).

11. Averina, O.A. et al. Mitochondrial peptide Mtln contributes to oxidative metabolism in mice. Biochimie 204, 136–139 (2023).

12. Friesen, M. et al. Mitoregulin Controls β-Oxidation in Human and Mouse Adipocytes. Stem Cell Reports 14, 590–602 (2020).

13. Averina, O.A. et al. Mitoregulin Contributes to Creatine Shuttling and Cardiolipin Protection in Mice Muscle. Int J Mol Sci 24 (2023).

14. Zhang, S. et al. LINC00116-encoded microprotein mitoregulin regulates fatty acid metabolism at the mitochondrial outer membrane. iScience 26, 107558 (2023).

15. Xiao, M.H. et al. Downregulation of a mitochondrial micropeptide, MPM, promotes hepatoma metastasis by enhancing mitochondrial complex I activity. Mol Ther 30, 714–725 (2022).

16. Wang, L. et al. The micropeptide LEMP plays an evolutionarily conserved role in myogenesis. Cell Death Dis 11, 357 (2020).

17. Averina, O.A. et al. Kidney-Related Function of Mitochondrial Protein Mitoregulin. Int J Mol Sci 24 (2023).

18. Li, J. et al. Mitochondrial micropeptide MOXI promotes fibrotic gene transcription by translocation to the nucleus and bridging N-acetyltransferase 14 with transcription factor c-Jun. Kidney Int 103, 886–902 (2023).

19. Davidson, S.M. et al. Mitochondrial and mitochondrial-independent pathways of myocardial cell death during ischaemia and reperfusion injury. J Cell Mol Med 24, 3795–3806 (2020).

20. Del Re, D.P., Amgalan, D., Linkermann, A., Liu, Q. & Kitsis, R.N. Fundamental Mechanisms of Regulated Cell Death and Implications for Heart Disease. Physiol Rev 99, 1765–1817 (2019).

21. Kwong, J.Q. & Molkentin, J.D. Physiological and pathological roles of the mitochondrial permeability transition pore in the heart. Cell Metab 21, 206–214 (2015).

22. Baines, C.P. et al. Loss of cyclophilin D reveals a critical role for mitochondrial permeability transition in cell death. Nature 434, 658–662 (2005).

23. Jang, S. et al. Elucidating Mitochondrial Electron Transport Chain Supercomplexes in the Heart During Ischemia-Reperfusion. Antioxid Redox Signal 27, 57–69 (2017).

24. Bernardi, P. et al. Identity, structure, and function of the mitochondrial permeability transition pore: controversies, consensus, recent advances, and future directions. Cell Death Differ 30, 1869–1885 (2023).

25. Karch, J. et al. Inhibition of mitochondrial permeability transition by deletion of the ANT family and CypD. Sci Adv 5, eaaw4597 (2019).

26. Urbani, A. et al. Purified F-ATP synthase forms a Ca(2+)-dependent high-conductance channel matching the mitochondrial permeability transition pore. Nat Commun 10, 4341 (2019).

27. Endlicher, R., Drahota, Z., Štefková, K., Červinková, Z. & Kučera, O. The Mitochondrial Permeability Transition Pore-Current Knowledge of Its Structure, Function, and Regulation, and Optimized Methods for Evaluating Its Functional State. Cells 12 (2023).

28. Carroll, K.J. et al. A mouse model for adult cardiac-specific gene deletion with CRISPR/Cas9. Proc Natl Acad Sci U S A 113, 338–343 (2016).

29. Mitchell, W. et al. The mitochondria-targeted peptide SS-31 binds lipid bilayers and modulates surface electrostatics as a key component of its mechanism of action. J Biol Chem 295, 7452–7469 (2020).

30. Pennington, E.R. et al. Proteolipid domains form in biomimetic and cardiac mitochondrial vesicles and are regulated by cardiolipin concentration but not monolyso-cardiolipin. J Biol Chem 293, 15933–15946 (2018).

31. Zoeller, R.A., Morand, O.H. & Raetz, C.R. A possible role for plasmalogens in protecting animal cells against photosensitized killing. J Biol Chem 263, 11590–11596 (1988).

32. Reiss, D., Beyer, K. & Engelmann, B. Delayed oxidative degradation of polyunsaturated diacyl phospholipids in the presence of plasmalogen phospholipids in vitro. Biochem J 323 (Pt 3), 807–814 (1997).

33. Kulik, W. et al. Bloodspot assay using HPLC-tandem mass spectrometry for detection of Barth syndrome. Clin Chem 54, 371–378 (2008).

34. Allen, M.E. et al. The cardiolipin-binding peptide elamipretide mitigates fragmentation of cristae networks following cardiac ischemia reperfusion in rats. Commun Biol 3, 389 (2020).

35. Acin-Perez, R. et al. A novel approach to measure mitochondrial respiration in frozen biological samples. Embo j 39, e104073 (2020).

36. Shimada, K. & Asahina, E. Visualization of intracellular ice crystals formed in very rapidly frozen cells at -27 degree C. Cryobiology 12, 209–218 (1975).

37. Karlsson, J.O. & Toner, M. Long-term storage of tissues by cryopreservation: critical issues. Biomaterials 17, 243–256 (1996).

38. Sherman, J.K. Correlation in ultrastructural cryoinjury of mitochondria with aspects of their respiratory function. Exp Cell Res 66, 378–384 (1971).

39. Štětina, T., Des Marteaux, L.E. & Koštál, V. Insect mitochondria as targets of freezing-induced injury. Proc Biol Sci 287, 20201273 (2020).

40. Quinn, P.J. A lipid-phase separation model of low-temperature damage to biological membranes. Cryobiology 22, 128–146 (1985).

41. Drobnis, E.Z. et al. Cold shock damage is due to lipid phase transitions in cell membranes: a demonstration using sperm as a model. J Exp Zool 265, 432–437 (1993).

42. Burger, K.N. Greasing membrane fusion and fission machineries. Traffic 1, 605–613 (2000).

43. Stepanyants, N. et al. Cardiolipin’s propensity for phase transition and its reorganization by dynamin-related protein 1 form a basis for mitochondrial membrane fission. Mol Biol Cell 26, 3104–3116 (2015).

44. Jouhet, J. Importance of the hexagonal lipid phase in biological membrane organization. Front Plant Sci 4, 494 (2013).

45. Rand, R.P. & Sengupta, S. Cardiolipin forms hexagonal structures with divalent cations. Biochim Biophys Acta 255, 484–492 (1972).

46. Pennington, E.R., Funai, K., Brown, D.A. & Shaikh, S.R. The role of cardiolipin concentration and acyl chain composition on mitochondrial inner membrane molecular organization and function. Biochim Biophys Acta Mol Cell Biol Lipids 1864, 1039–1052 (2019).

47. Smaal, E.B. et al. Calcium-induced changes in permeability of dioleoylphosphatidylcholine model membranes containing bovine heart cardiolipin. Biochim Biophys Acta 897, 191–196 (1987).

48. Ortiz, A., Killian, J.A., Verkleij, A.J. & Wilschut, J. Membrane fusion and the lamellar-to-inverted-hexagonal phase transition in cardiolipin vesicle systems induced by divalent cations. Biophys J 77, 2003–2014 (1999).

49. Fox, C.A., Ellison, P., Ikon, N. & Ryan, R.O. Calcium-induced transformation of cardiolipin nanodisks. Biochim Biophys Acta Biomembr 1861, 1030–1036 (2019).

50. Gohrbandt, M. et al. Low membrane fluidity triggers lipid phase separation and protein segregation in living bacteria. Embo j 41, e109800 (2022).

51. Eble, K.S., Coleman, W.B., Hantgan, R.R. & Cunningham, C.C. Tightly associated cardiolipin in the bovine heart mitochondrial ATP synthase as analyzed by 31P nuclear magnetic resonance spectroscopy. J Biol Chem 265, 19434–19440 (1990).

52. Hoffmann, B., Stöckl, A., Schlame, M., Beyer, K. & Klingenberg, M. The reconstituted ADP/ATP carrier activity has an absolute requirement for cardiolipin as shown in cysteine mutants. J Biol Chem 269, 1940–1944 (1994).

53. Paradies, G., Paradies, V., Ruggiero, F.M. & Petrosillo, G. Role of Cardiolipin in Mitochondrial Function and Dynamics in Health and Disease: Molecular and Pharmacological Aspects. Cells 8 (2019).

54. Bround, M.J., Bers, D.M. & Molkentin, J.D. A 20/20 view of ANT function in mitochondrial biology and necrotic cell death. J Mol Cell Cardiol 144, A3–a13 (2020).

55. Miranda É, G.A. et al. Cardiolipin Structure and Oxidation Are Affected by Ca(2+) at the Interface of Lipid Bilayers. Front Chem 7, 930 (2019).

56. Paradies, G. et al. Decrease in mitochondrial complex I activity in ischemic/reperfused rat heart: involvement of reactive oxygen species and cardiolipin. Circ Res 94, 53–59 (2004).

57. Zhao, K. et al. Cell-permeable peptide antioxidants targeted to inner mitochondrial membrane inhibit mitochondrial swelling, oxidative cell death, and reperfusion injury. J Biol Chem 279, 34682–34690 (2004).

58. Birk, A.V. et al. The mitochondrial-targeted compound SS-31 re-energizes ischemic mitochondria by interacting with cardiolipin. J Am Soc Nephrol 24, 1250–1261 (2013).

59. Dai, D.F. et al. Mitochondrial targeted antioxidant Peptide ameliorates hypertensive cardiomyopathy. J Am Coll Cardiol 58, 73–82 (2011).

60. Hou, Y. et al. Mitochondria-targeted peptide SS-31 attenuates renal injury via an antioxidant effect in diabetic nephropathy. Am J Physiol Renal Physiol 310, F547–559 (2016).

61. Chiao, Y.A. et al. Late-life restoration of mitochondrial function reverses cardiac dysfunction in old mice. Elife 9 (2020).

62. Bomba-Warczak, E., Edassery, S.L., Hark, T.J. & Savas, J.N. Long-lived mitochondrial cristae proteins in mouse heart and brain. J Cell Biol 220 (2021).

63. Xu, Y. et al. Loss of protein association causes cardiolipin degradation in Barth syndrome. Nat Chem Biol 12, 641–647 (2016).

64. Moon, S.H. et al. Activation of mitochondrial calcium-independent phospholipase A2γ (iPLA2γ) by divalent cations mediating arachidonate release and production of downstream eicosanoids. J Biol Chem 287, 14880–14895 (2012).

65. Jabůrek, M., Průchová, P., Holendová, B., Galkin, A. & Ježek, P. Antioxidant Synergy of Mitochondrial Phospholipase PNPLA8/iPLA2γ with Fatty Acid-Conducting SLC25 Gene Family Transporters. Antioxidants (Basel) 10 (2021).

66. Birk, A.V., Chao, W.M., Liu, S., Soong, Y. & Szeto, H.H. Disruption of cytochrome c heme coordination is responsible for mitochondrial injury during ischemia. Biochim Biophys Acta 1847, 1075–1084 (2015).

67. Kagan, V.E. et al. Anomalous peroxidase activity of cytochrome c is the primary pathogenic target in Barth syndrome. Nat Metab 5, 2184–2205 (2023).

68. Kutschka, I. et al. Activation of the integrated stress response rewires cardiac metabolism in Barth syndrome. Basic Res Cardiol 118, 47 (2023).

69. Díaz, F., Barrientos, A. & Fontanesi, F. Evaluation of the mitochondrial respiratory chain and oxidative phosphorylation system using blue native gel electrophoresis. Curr Protoc Hum Genet Chapter 19, Unit19.14 (2009).

70. Jha, P., Wang, X. & Auwerx, J. Analysis of Mitochondrial Respiratory Chain Supercomplexes Using Blue Native Polyacrylamide Gel Electrophoresis (BN-PAGE). Curr Protoc Mouse Biol 6, 1–14 (2016).

71. Pennington, E.R. et al. Distinct membrane properties are differentially influenced by cardiolipin content and acyl chain composition in biomimetic membranes. Biochim Biophys Acta Biomembr 1859, 257–267 (2017).

72. Matyash, V., Liebisch, G., Kurzchalia, T.V., Shevchenko, A. & Schwudke, D. Lipid extraction by methyl-tert-butyl ether for high-throughput lipidomics. J Lipid Res 49, 1137–1146 (2008).

73. Molenaar, M.R. et al. LION/web: a web-based ontology enrichment tool for lipidomic data analysis. Gigascience 8 (2019).

